# Myeloma and therapy reshape the bone marrow niche to durably constrain immune reconstitution and vaccine responsiveness

**DOI:** 10.64898/2026.04.08.717307

**Authors:** Aishwarya Chander, Samir Rachid Zaim, Medbh A. Dillon, Palak C. Genge, Nicholas Moss, Patrick I. McGrath, Mackenzie S. Kopp, Kevin J. Lee, Emma L. Kuan, Julian Reading, Veronica Hernandez, Xiaoling Song, Mansi Singh, Jessica Garber, Christian M. LaFrance, Garth L. Kong, Marla C. Glass, Eden Leigh W. Davis, David Glass, Yudong D. He, Alexander T. Heubeck, Erin K. Kawelo, Upaasana Krishnan, Cara Lord, Paul Meijer, Regina R. Mettey, Blessing Musgrove, Lauren Y. Okada, Vaishnavi Parthasarathy, Tao Peng, Cole G. Phalen, Stanley Riddell, Charles R. Roll, Tyanna J. Stuckey, Elliott G. Swanson, Zach J. Thomson, Morgan D.A. Weiss, Peter J. Wittig, Stephanie D. Anover-Sombke, Ernest M. Coffey, Lynne A. Becker, Thomas F. Bumol, Ananda W. Goldrath, Evan W. Newell, Philip D. Greenberg, Xiao-jun Li, Susan M. Kaech, Peter J. Skene, Lucas T. Graybuck, Love Tätting, Mikael Sigvardsson, Mary Kwok, Damian J. Green, Troy R. Torgerson, Melinda L. Angus-Hill

## Abstract

Infections are the most common cause of non-relapse mortality in multiple myeloma (MM), but the basis of persistent immune dysfunction is obscured by patient heterogeneity and complex treatment regimens, including autologous stem cell transplant (ASCT). We performed longitudinal multi-omic profiling of matched bone marrow and peripheral blood from MM patients across diagnosis, induction, ASCT, and recovery. We found the tumor imposes a compartment-specific immune program where the marrow exhibits metabolic and inflammatory changes that bias hematopoiesis and alter cytotoxic effector programs not mirrored in blood. Adaptive immune reconstitution is impaired up to two years post-ASCT. Half of patients fail to mount IgG responses to high-dose non-adjuvanted influenza vaccine, a defect overcome by the lipid nanoparticle (LNP) adjuvanted COVID mRNA vaccine, which elicited responses in all patients, supporting adjuvanted influenza vaccine strategies in MM. Together these findings define how myeloma and its treatment durably reshape immunity from the marrow outward.

**Highlights:**

- Multiple Myeloma marrow and blood show opposing metabolic and inflammatory states
- Induction therapy selects durable myeloma plasma-cell transcriptional states
- B cell and follicular helper T deficits blunt antigen responses after transplant
- COVID-19 vaccination builds immune memory with variable responses to flu vaccination

**eTOC:** Multiple myeloma and its treatment leave a lasting imprint on the bone marrow niche. By profiling bone marrow and blood longitudinally at diagnosis, through induction, autologous transplant, and recovery, we show that marrow-local metabolic and inflammatory constraints persist and help explain why influenza vaccination often fails while mRNA vaccination succeeds.

## Introduction

Multiple myeloma (MM) is a bone marrow resident plasma cell malignancy that causes complex clinical manifestations including hypercalcemia, renal dysfunction, anemia and osteolytic bone lesions (CRAB)^1^. Although generally considered chronic and largely incurable, modern therapy aims to achieve deep remission to prevent organ damage and prolong survival. In transplant-eligible patients, standard induction therapy combines an immunomodulatory drug, a proteasome inhibitor, and a corticosteroid drug, often with an anti-CD38 antibody to deepen response^2,3^. Although outcomes have improved, relapse and clinically significant infections remain frequent.

Infection is a major cause of mortality^4^, underscoring the importance of disease- and treatment-associated immune dysfunction. Approximately 70% of newly diagnosed multiple myeloma patients face clinically significant infections, with infection-related mortality reaching 27% within the first year after diagnosis^4^. Infection risk is elevated, not only after diagnosis and treatment, but also in the months preceding diagnosis, and can remain high for several years^4,5^. However, we do not know which immune and hematopoietic abnormalities arise at diagnosis, how they evolve through induction therapy and transplant, or which deficits fail to resolve during immune reconstitution. A central challenge in studying MM is its compartmentalization, the bone marrow is both the site of disease and hematopoiesis, yet most immune monitoring is performed in blood. Recent single-cell profiling of long-term survivors suggests that marrow immune alterations can persist for many years after first-line therapy, raising the possibility of durable immune “scarring”. When these alterations arise across induction and transplant and how they relate to concurrent blood changes remains unclear^6^.

We therefore asked three key questions. First, which tumor cell states persist despite standard therapy, and whether transcriptional state provides information about persistence after induction that is not captured by cytogenetic features alone. Second, which immune and hematopoietic abnormalities are present at diagnosis, how they are reshaped by induction therapy and transplant, and whether immune recovery in the bone marrow mirrors what is observed in the blood. Third, after therapy, do persistent abnormalities reflect continued loss of specific lineages or lasting functional changes within reconstituted cells, and do these defects help explain impaired vaccine responses following transplant?

To address these questions, we performed prospective longitudinal multi-omic profiling of newly diagnosed multiple myeloma patients (NDMM) at diagnosis and throughout treatment with bortezomib, lenalidomide, and dexamethasone (VRd) induction therapy, followed by consolidation and autologous stem cell transplant (ASCT). We profiled matched bone marrow and peripheral blood using single-cell RNA sequencing (scRNA-seq), complemented by flow cytometry and proteomic profiling of plasma and bone marrow interstitial fluid, and integrated these assays with post-transplant vaccine responses to quantify the restoration of functional immunity. Together, these data define how therapy reshapes immune and hematopoietic compartments and point to vaccination strategies to improve protection from infection after transplant. The results are organized around therapy-selected tumor states that persist after induction, compartment-specific immune remodeling in marrow versus blood, and progenitor and lymphocyte reconstitution linked to functional vaccine outcomes. We also provide tools to explore the multi-modal datasets from this manuscript using an interactive viewer https://apps.allenimmunology.org/aifi/insights/ndmm/.

## Results

### A representative NDMM cohort profiled longitudinally across VRd and ASCT

We used a multi-omic systems immunology approach to profile matched bone marrow and peripheral blood longitudinally and define the cellular and molecular effects of myeloma and standard-of-care therapy (VRd + ASCT) on the immune system and microenvironment. We enrolled 17 transplant-eligible newly diagnosed multiple myeloma patients treated with VRd induction followed by high-dose melphalan (HDM) and ASCT, and collected paired marrow and blood from diagnosis through induction and post-transplant recovery (**Fig. 1A**; **Table 1**). Samples were obtained at diagnosis (N = 17), during induction (N = 14), after transplant (N = 14), and during maintenance at 1 year (N = 13) and 2 years (N = 7) (**Fig. 1A**). Baseline demographics and RISS distribution were comparable to published NDMM cohorts (**Table 1**; **Fig. 1B**)^7–9^. By study end, 66.7% achieved complete or stringent complete response and 60% improved response over therapy, with 13.3% progressing (**Fig. 1B; Supp. Table 1A**), supporting the generalizability of this cohort for longitudinal immune and microenvironmental analyses. A graphical representation of the study design and the total number of samples and cells or analytes evaluated are shown in **Fig. 1A** and **Fig. 1C**, respectively. Matched longitudinal sampling enabled separation of compartment-specific features from systemic signals and distinction of diagnosis-associated effects from treatment-driven reconstitution.

**Figure 1:**
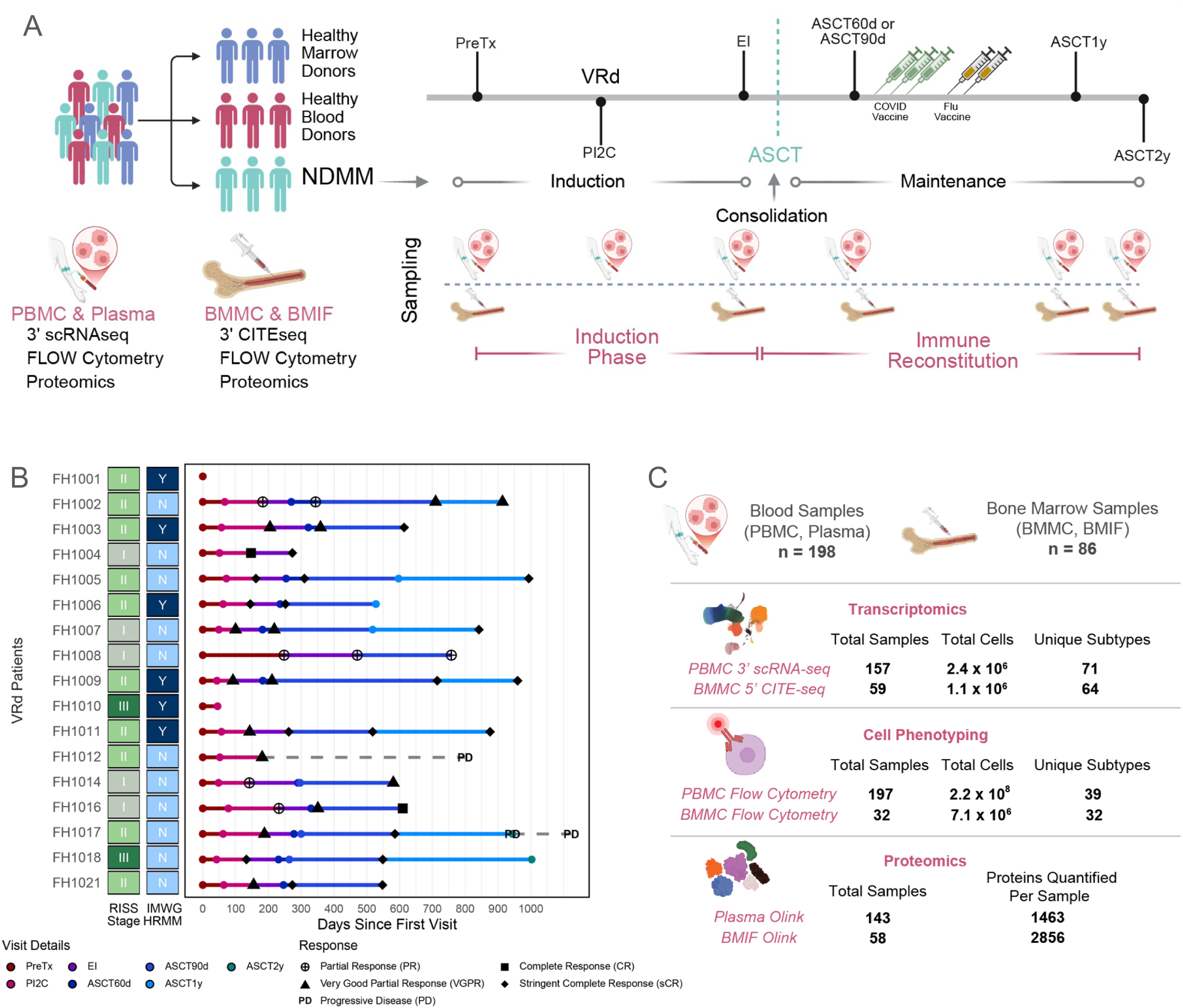
Study design, longitudinal sampling, and cohort overview. **(A)** Overview and sampling schema of the VRd treated NDMM cohort of multiple myeloma patients (n=17), healthy blood donors (n=32), and healthy bone marrow donors (n=10). Sampling schematic for the VRd-treated NDMM cohort with paired PBMC/plasma and BMMC/BMIF collected across treatment phases and timepoints; assays per compartment shown. **(B)** Per-patient sampling timeline (n = 17) showing days since first visit, RISS stage, IMWG HRMM status, and clinician-recorded response at each timepoint. **(C)** Sample counts and analyte yield across scRNA-seq, flow cytometry, and Olink proteomics for blood and bone marrow compartments. **Abbreviations:** Newly diagnosed multiple myeloma (NDMM), bortezomib/lenalidomide/dexamethasone (VRd), peripheral blood mononuclear cells (PBMC), bone marrow mononuclear cells (BMMC), bone marrow interstitial fluid (BMIF), Pre-Treatment (PreTx), Post-Induction 2 Cycles (PI2C), End of Induction (EI), autologous stem cell transplant (ASCT), 60 and 90 Days Post-Transplant (ASCT60d, ASCT90d), 1 and 2 Year Post-Transplant (ASCT1y, ASCT2y), Revised International Staging System (RISS), International Myeloma Working Group high-risk multiple myeloma (IMWG HRMM).

**Table 1:** NDMM cohort demographics.

| Variable | Category | VRd (n=17) |
| --- | --- | --- |
| Sex |  |  |
|  | Female | 9 (52.9%) |
|  | Male | 8 (47.1%) |
| Race |  |  |
|  | Other | 2 (11.8%) |
|  | African American | 1 (5.9%) |
|  | Caucasian | 14 (82.4%) |
| Ethnicity |  |  |
|  | Hispanic or Latino origin | 0 (0.0%) |
|  | Non-Hispanic origin | 15 (88.2%) |
| Other |  | 2 (11.8%) |
| ageAtEnrollment |  |  |
| | Mean (SD) | 59.5 (± 9.0) |
|  | Median [IQR] | 62.0 [58.0, 65.0] |
|  | Range | 33.0–70.0 |
| RISS.stage |  |  |
|  | I | 5 (29.4%) |
|  | II | 10 (58.8%) |
|  | III | 2 (11.8%) |

### Malignant plasma cell states span cytogenetic backgrounds and show state-selective persistence after induction

To capture heterogeneous tumor effects on the immune system, we combined BMMC scRNA-seq data from 17 MM patients and 10 healthy donors into a single dataset spanning five time points (**Fig. 2A**). Projection of plasma cells onto a 2D Uniform Manifold Approximation and Projection (UMAP) revealed distinct patient-to-patient transcriptional heterogeneity, whereas healthy plasma cells showed more homogeneity (**Fig. 2B**). Because malignant transcriptional heterogeneity can reflect underlying genomic structure, we asked whether inferred copy number variations (CNVs) account for the shared and patient-specific state organization^10–12^. Clinical tumor cytogenetics, when available, together with **in silico** CNV inference using inferCNV, corroborated known cytogenetic features and provided additional CNV predictions, indicating that CNV heterogeneity is present and measurable in this cohort, including prediction of 1p loss in confirmed cases and detection of MAFB overexpression consistent with the effects of t14:20 translocation (**Supp. Table 2; Supp. Fig. 1A-B)**.

**Figure 2:**
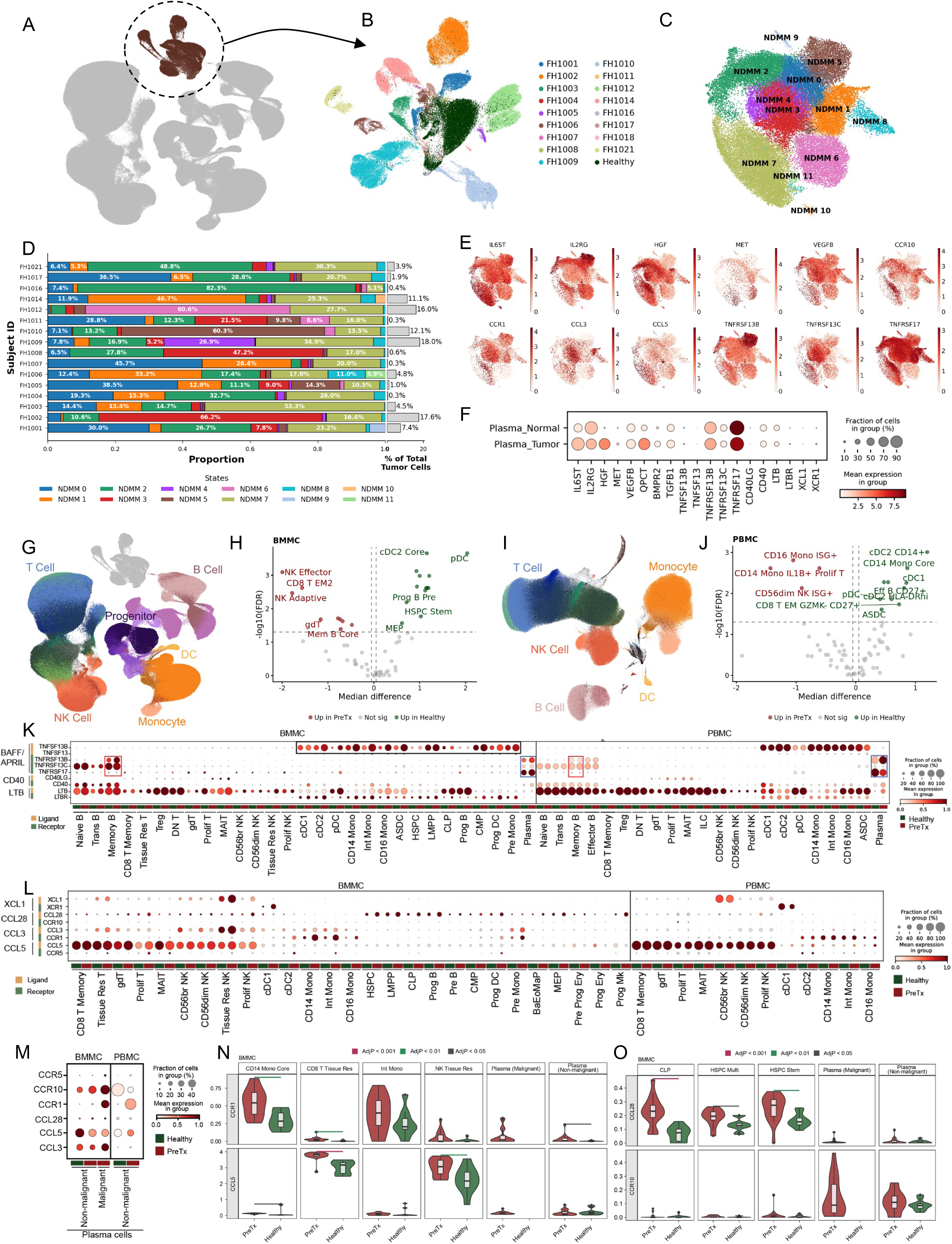
Malignant plasma cells and effects on the immune compartment. **(A)** UMAP of BMMC scRNA-seq at PreTx from the VRd-treated NDMM cohort, highlighting malignant and non-malignant plasma cells. **(B)** UMAP of plasma cells including PreTx malignant, PreTx non-malignant, and external healthy subsets, colored by donor. **(C)** Malignant plasma cell UMAP colored by integrated Leiden states (NDMM_0-11). **(D)** Per-patient distribution of malignant plasma cells across transcriptional states, colored by state at PreTx, with proportion of tumor cells per cluster and percent of represented tumor cells per patient. **(E-F)** Expression of selected immune reactive genes across plasma cell states (feature plots) and in malignant versus non-malignant plasma cells (dot plot). **(G, I)** UMAPs of immune subsets exclusive of plasma cells in BMMC **(G)** and PBMC **(I)** at PreTx with major lineages annotated. **(H, J)** Differential abundance of scRNA-seq cell subsets in PreTx versus healthy donors for BMMC **(H)** and PBMC **(J)**, summarized as median CLR difference and -log10FDR. **(K-M)** Ligand-receptor axes across BMMC and PBMC subsets, including lymphogenesis **(K)** and chemokine/trafficking axes **(L)**, with a malignant and healthy plasma cell-focus in PBMC and BMMC **(M)**. **(N-O)** Pseudobulk expression in selected BMMC subsets comparing PreTx and healthy donors. Horizontal bars indicate significance; colors denote AdjP. Abbreviations: centered log-ratio (CLR); false discovery rate (FDR).

To determine whether malignant transcriptional states map to cytogenetic background, we integrated malignant plasma cells across patients in pre-treatment samples (PreTx) using Harmony on the PCA embedding and identified clusters using the Leiden algorithm^13,14^. This analysis identified nine shared malignant plasma cell states (NDMM_0-8) and three patient-specific clusters (9-11; **Fig. 2C-D**; **Supp. Fig. 1C-E**). We focused on the shared states and aligned them to published MM expression signatures^15,16^, revealing interferon-responsive features in NDMM_4, proliferative features in NDMM_8, and monocyte-like features in NDMM_7^17^ (**Supp. Fig. 1F-G; Supp. Table 6K**). NDMM_5 co-enriched for high risk features including *CXCR4*, *MMSET/FGFR3*, *MAF*, and *APOBEC* (**Supp. Fig. 1G**)^18^. Each patient contributed cells to multiple transcriptional states irrespective of genetic background (**Fig. 2D; Supp. Fig. 1C-E**).

We next performed gene set enrichment analysis (GSEA) using FGSEA to identify signatures that distinguish malignant plasma cell states and could possibly shape immune interactions. Metabolic, signaling, and chromatin regulatory gene sets varied by state relative to healthy plasma cells, including enrichment of a SUZ12 ENCODE signature in a subset of states (**Supp. Fig. 1H; Supp. Table 6Y**). MYC-associated pathways were inversely related to SUZ12 enrichment, consistent with coordinated changes in PRC2-linked repression and MYC-driven transcription^19–22^. Immune and inflammatory gene sets, including TNF-α signaling via NF-κB, were broadly attenuated across malignant states. Immunoactive genes highlighted some state-specific ligands and receptors, including BAFF axis components and growth factors such as *HGF*, *MET*, and *VEGFB* (**Fig. 2E-F**), and several of these proteins were detected in plasma proteomics, including HGF, with correlated expression between compartments (BMIF vs. plasma) at PreTx (**Supp. Fig. 1I-J**), while TNF showed poor correlation of expression.

After defining malignant plasma cell states at diagnosis, we next asked which malignant states persist after VRd induction, and whether induction sensitivity reflects transcriptional state (**Fig. 2C; Supp. Table 1A**). Following VRd, clusters NDMM_1, NDMM_4, NDMM_6 and patient-specific clusters NDMM_9-11 were largely lost (**Supp. Fig. 1K**). NDMM_4 and NDMM_6 were positive for inflammatory and protein synthesis signatures (**Supp. Fig. 1G-H**), and NDMM_1 showed increased ribosome biogenesis signatures (NES = 1.92; padj = 0.023; **Supp. Table 6Y**), and all three showed enrichment for the SMM-8 patient specific cluster characterized by proteostatic burden ^15,16^. Together these findings indicate that multiple VRd-sensitive states converge on translational activation, ribosome biogenesis and proteostatic stress (**Supp. Fig. 1G**). In contrast, despite cytogenetically dominated transcriptional states such as NDMM_3, several other malignant states persisted, including NDMM_0, NDMM_2, NDMM_5, NDMM_7, and NDMM_8 (**Supp. Fig. 1K**). Notably, although NDMM_5 was rare at diagnosis, its fractional contribution increased across all patients following therapy (**Fig. 2D; Supp. Fig. 1K**). Persistent states spanned oxidative phosphorylation-associated states (NDMM_0/2), *APOBEC*, *CXCR4*, and *MMSET/MAF* signatures (NDMM_5**; Supp. Fig. 1G**), and a proliferative state (NDMM_8**; Supp. Fig. 1H**). Taken together, these data suggest that VRd imposes strong selection against biosynthetically active tumor states, enriching for transcriptionally distinct residual states.

### At diagnosis, immune remodeling differs between marrow and blood

We profiled matched marrow and blood at diagnosis to define myeloma-associated remodeling of immune and hematopoietic progenitor composition prior to treatment. In bone marrow, we analyzed 69 samples from patients (N=17) and healthy donors (N=10) and identified 64 cell subsets (**Supp. Table 6**; **Fig. 2G**). Relative to healthy controls, myeloma marrow collected pre-treatment (BMMC PreTx) showed expansion of cytotoxic populations, including NK effector and CD8 effector memory (EM) subsets, increased frequencies of ISG⁺ naïve lymphocytes and select B cell subsets, and reductions in multiple progenitor and myeloid lineages (**Fig. 2H**). In PBMC, standardized cell labeling annotated 71 cell types in patients and 32 age-matched healthy donors (**Supp. Table 5**; **Fig. 2I**)^23^. NDMM PBMCs had altered B cell composition, modest shifts in cytotoxic and select CD8+ EM states, expansion of IL1B⁺ CD14 monocytes and reduced representation of other monocyte and dendritic cell (DC) populations (**Fig. 2J; Supp. Table 5**)^24,25^. Together, these data indicate myeloma-associated compartment specific immune remodeling, with marrow showing cytotoxic enrichment and disruption of progenitor and antigen-presenting subsets whereas blood shows dominant myeloid activation characterized by IL1β and IFN-associated monocyte states.

We analyzed B cell survival and proliferation signaling across blood and marrow (**Fig. 2K; Supp. Fig. 1J**). *BAFF* transcription (*TNFSF13B*) was increased in marrow myeloid and progenitor subsets without detectable change in peripheral blood, and BAFF protein levels were correlated between BMIF and plasma at PreTx (**Fig. 2K; Supp. Fig. 1J**). In contrast, *APRIL* (*TNFSF13*) remained low with no disease-associated change (**Fig. 2K**). BAFF receptor (*TNFRSF13B)* was elevated at PreTx, while APRIL receptor *TNFRSF13C* was reduced at PreTx in marrow-associated memory B Cells relative to healthy subsets (**Fig. 2K**). The CD40 axis showed compartment-specific modulation with increased *CD40* in PreTx marrow myeloid subsets, while the LTB-LTBR axis showed no clear disease-related changes (**Fig. 2K**). Together, these data indicate marrow-local tuning of B cell survival signaling at diagnosis.

We next examined chemokine and trafficking gene sets involved in homing, migration, and retention. At PreTx, these signatures were more perturbed in marrow than blood, with disease-associated increases in *CCR1* in monocytes and DCs and increased *CCR5* across multiple BMMC subsets (**Fig. 2L**). In marrow, *CCL3* was broadly expressed with disease-associated modulation, whereas *CCL5* was largely confined to cytotoxic lymphocytes, while PBMCs showed very low *CCL3* with more stable receptor patterns (**Fig. 2L**, **Fig. 2N-O**). Plasma cells showed the strongest compartmental contrast. In marrow, both PreTx non-malignant and malignant plasma cells had reduced CCL5, with malignant cells showing higher CCL3 and altered receptor usage (including CCR1 and CCR10). In blood, plasma cells lacked CCL3 and showed higher CCL5 at PreTx than healthy controls, and the dominant plasma-cell receptor signature shifted from CCR10 in healthy to CCR1 in PreTx non-malignant plasma cells (**Fig. 2L; BMMC in Fig. 2M–O**). Together, these findings show compartmentalized trafficking signatures with stronger marrow perturbation and plasma cell-specific differences across marrow and blood.

### Integrated flow, single-cell, and proteomic profiling maps global immune remodeling across therapy

To define how diagnosis-associated immune alterations evolve with treatment, we integrated longitudinal flow cytometry, scRNA-seq, and paired plasma and BMIF proteomics across induction and transplant. Flow cytometry and absolute lymphocyte count (ALC)-adjusted cell counts showed reduced circulating DCs and CD14 monocytes at diagnosis, with reductions in selected memory B cells, T follicular helper (Tfh) cells, and CD8 resident memory subsets, while most circulating B cell counts were similar to healthy controls (**Fig. 3A; Supp. Fig. 2A-B; Supp. Table 3**)^26^. By the end of induction (EI), several myeloid defects improved, whereas broader B cell lineage depletion emerged and CD4 and unconventional T cell populations declined (**Fig. 3A; Supp. Fig. 2B**)^27,28^. Separately, an additional exploratory cohort receiving daratumumab plus VRd (DVRd) showed expected CD38-lineage effects at EI (**Supp. Fig. 3B-D**), mostly in specific CD56 dim NK subsets^29,30^. In our primary VRd-only cohort, post-ASCT reconstitution was polarized, with persistent deficits in helper and central memory populations and expansion of differentiated effector states (**Fig. 3A; Supp. Fig. 3A**; **Fig. 6**).

**Figure 3:**
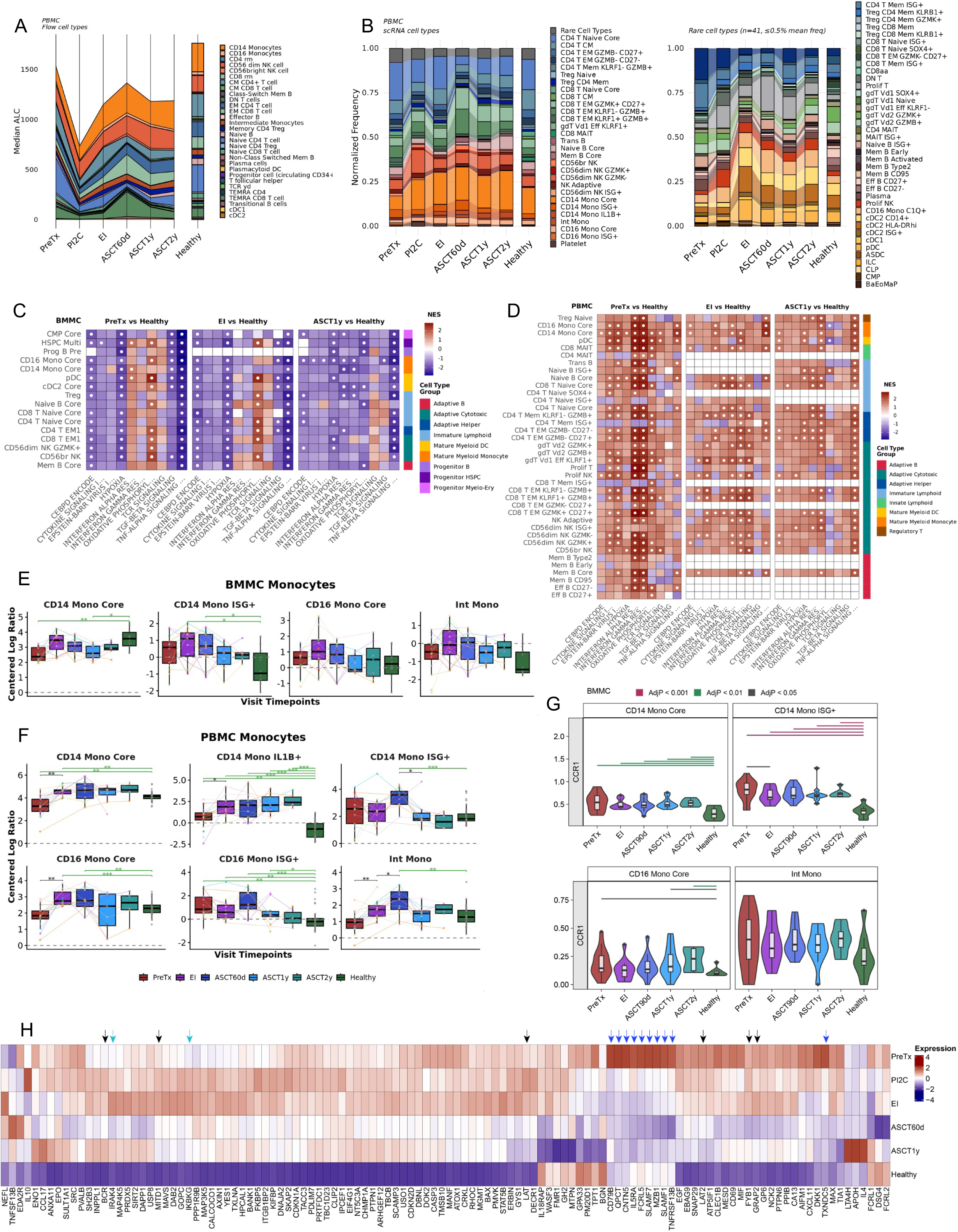
Global multi-assay overview of immune composition and pathway signatures across therapy. **(A)** Circulating immune cell median ALC from PBMC flow cytometry across treatment timepoints, shown alongside healthy donors. **(B)** scRNA-seq PBMC cell type frequencies across timepoints, with rare populations collapsed in the main view and shown separately. **(C-D)** Gene set enrichment (NES) comparing each indicated timepoint with healthy controls across representative cell types in BMMCs **(C)** and PBMCs **(D)**; white dots denote AdjP<0.05. **(E-F)** CLR-transformed monocyte subset frequencies across timepoints in BMMCs **(E)** and PBMCs **(F)** relative to healthy controls. **(G)** Pseudobulk gene expression in BMMC monocyte states across timepoints and healthy controls. **(H)** Heatmap showing relative expression of differential plasma proteins across timepoints relative to healthy controls. Arrows denote selected proteins, grouped by functional class (TCR/BCR, black; tumor-related, blue; and NF-κB signaling, cyan). Abbreviations: absolute lymphocyte counts (ALC), centered log-ratio (CLR), normalized enrichment score (NES).

Using transcriptionally defined states, scRNA-seq compositional shifts in PBMC broadly mirrored the flow cytometry patterns across therapy (**Fig. 3B**; **Supp. Fig. 2E,G,I; Supp. Table 5**). In marrow, scRNA-seq revealed persistent reductions in DC, monocyte and progenitor subsets at PreTx alongside cytotoxic EM and IFN-skewed immune states and increases in some B cell populations (**Supp. Fig. 2C,D,F,H; Supp. Table 6**). By EI, depletion across pDC, progenitor and select B cell subsets persisted, while inflammatory and cytotoxic subsets remained enriched. Marrow at 90 days after ASCT (ASCT90d) showed marked depletion of progenitor, antigen-presenting subsets, alongside reductions in adaptive populations with reciprocal enrichment of cytotoxic and innate states and B cell lineage precursor and transitional states (**Supp. Fig. 2F; Supp. Table 6**). By 1 or 2 years post ASCT (ASCT1y/ASCT2y), reconstitution remained incomplete with persistent dendritic and progenitor deficits and durable enrichment of select cytotoxic and B cell lineage progenitor states (**Supp. Fig. 2H; Supp. Table 6**).

Pathway enrichment results diverged between marrow and blood. In marrow, hypoxia signatures were repressed at PreTx and remained suppressed through ASCT1y across multiple lineages (**Fig. 3C**). Downstream of hypoxia regulation, the CEBPD ENCODE pathway was reduced across marrow subsets, whereas PBMCs showed a subset-specific increase by ASCT1y (**Fig. 3C-D**)^31^. TNF-α signaling via NF-κB tracked with hypoxia and remained attenuated across marrow subsets (**Fig. 3C; Supp. Table 6**)^32,33^, while both signatures were increased across many PBMC subsets (**Fig. 3D; Supp. Table 6**). Interferon response signatures also diverged, with broad elevation across PBMC subsets and more limited changes in marrow (**Fig. 3C-D**). TGFB1 was correlated between BMIF and plasma at PreTx, whereas TNF was not (**Supp. 1J**), but despite TGFB1 concordance, TGF-β pathway remained reduced in marrow and was not significantly altered in PBMCs (**Fig. 3C-D**), suggesting that ligand availability does not necessarily reflect downstream pathway activation across compartments. Taken together these results are consistent with strong compartmentalization of inflammatory signaling throughout treatment.

We next examined remodeling of three CD14⁺ monocyte states (core, ISG⁺, and IL1B⁺) across therapy in blood and marrow (**Fig. 3E-F**)^34^. In blood, IL1B⁺ monocytes were expanded at PreTx and persisted through ASCT2y. Core monocytes were reduced at PreTx but normalized by EI, while ISG⁺ monocytes were near healthy levels at diagnosis but peaked post transplant before returning to healthy levels by ASCT1y (**Fig. 3E-F**). In marrow, core monocytes recovered by EI, and were comparable to healthy post-transplant, while ISG⁺ monocytes showed a transient induction-associated increase before normalizing post-transplant (**Fig. 3E-F**). The IL1B⁺ population observed in blood was not identified in the BMMC dataset, potentially reflecting a shift toward tissue-resident macrophages over monocytes in the marrow environment (**Fig. 3E-F**). Building on *CCR1* expression changes (**Fig. 2N**), we found *CCR1* expression was highest in marrow CD14⁺ ISG⁺ monocytes and remained high across therapy relative to healthy controls (**Fig. 3G**). This marrow *CCR1* enrichment is consistent with increased *CCL3* and *CCL5* ligand availability (**Fig. 2L**)^35,36^. In contrast, in PBMCs, *CCR1* showed only modest variation across monocyte states and timepoints (**Supp. Fig. 2M**).

We next profiled matched BMIF and plasma proteomics longitudinally and relative to healthy controls. At PreTx, plasma proteomics was dominated by proteins linked to B and plasma cell biology and myeloma disease burden (**Fig. 3H**, blue arrows**; Supp. Fig. 2J,L; Supp. Table 4**), including FCRL5, TXNDC5, and SLAMF7^37,38^. PreTx to EI changes were concordant between plasma and BMIF (**Supp. Fig. 2L**), with decreases in multiple B lineage associated markers in both compartments (e.g., TCL1A, FCRL1, FCRL2), consistent with contraction of the malignant plasma cells and reduced B lineage signals following VRd therapy (**Fig. 3A-B; Supp. Fig. 2A-B**). In contrast, EI plasma showed increased abundance of T Cell Receptor (TCR) and B cell receptor (BCR) adaptors proteins, NF-κB associated signaling components, and BAFF (TNFSF13B; **Fig. 3H**, black and cyan arrows). Pathway enrichment analysis of differential plasma proteomics overlapped with PBMC FGSEA results, showing strongest enrichment of inflammatory pathways, including TNF-α signaling via NF-κB, fewer enriched pathways at EI, and re-emergence of several pathways post-transplant before normalizing by ASCT2y (**Supp. Fig. 2K**).

### Therapy and transplant reshape progenitor composition, metabolism and inflammatory pathway signatures

Because durable immune deficits may reflect constraints in hematopoietic output from the marrow niche, we quantified progenitor composition across therapy relative to healthy controls (**Fig. 4A; Supp. Fig. 4A**). We identified three compositional trajectories among progenitor populations: (i) **persistent depletion** of early HSPC and lymphoid-entry states (HSPC Stem/Multi, LMPP, CLP), (ii) **transient rebound** at EI or ASCT90d followed by decline in myeloid and DC progenitors (CMP Core and Prog DC cDC/pDC), and (iii) **progressive expansion** of committed B-lineage progenitors during recovery (**Fig. 4A; Supp. Fig. 4A**). The concurrent depletion of CLPs and expansion of committed B-lineage progenitors is consistent with rapid transit from CLP into Prog B Pre states, rather than an overall block in lymphoid lineage commitment. Overlaying pathway signatures showed that, despite these divergent compositional outcomes, progenitors shared a marrow-imposed transcriptional state characterized by broad suppression of hypoxia-associated gene sets and HIF1A ENCODE signatures, with concordant changes in nutrient sensing components (HIF1A down, LAMTOR1 up) (**Fig. 4B**; **Fig. 4C; Supp. Table 6**). Inflammatory pathway remodeling normalized more slowly than frequencies, with persistent disruption of TNF-α signaling via NF-κB that showed gradual recovery in leading edge genes over therapy (**Fig. 4B**; **Fig. 4D; Supp. Fig. 4E-F**)^39,40^.

**Figure 4:**
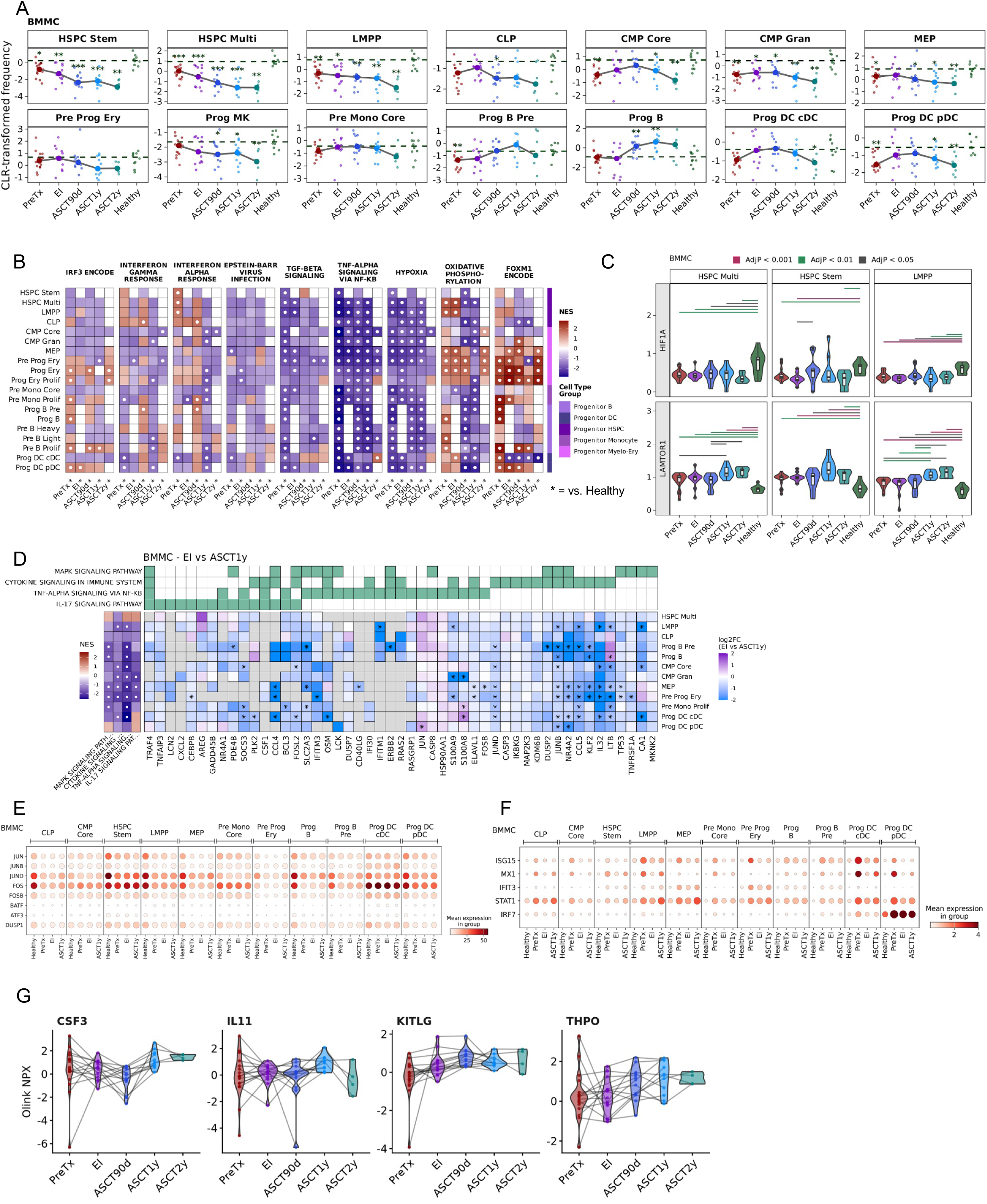
Hematopoietic progenitor subset dynamics and pathway remodeling across treatment. For BMMC subsets: **(A)** CLR-transformed frequencies of progenitor subsets across timepoints relative to healthy donors (green asterisks=AdjP versus healthy). **(B)** FGSEA pathway enrichment (NES) across progenitor subsets at each timepoint relative to healthy controls. **(C)** pseudobulk expression across timepoints. **(D)** EI to ASCT1y remodeling of inflammatory and signaling programs, showing pathway-level NES and leading-edge gene log2FC for the indicated pathways. **(E-F)** Dot plots of immediate-early/AP-1 genes **(E)** and interferon-stimulated genes **(F)** across progenitor subsets and timepoints. **(G)** BMIF proteomics of niche growth factors across timepoints. White dots and black-asterisks denote AdjP<0.05. Abbreviations: hematopoietic stem and progenitor cells (HSPC), lymphoid-primed multipotent progenitor (LMPP), common lymphoid progenitor (CLP), common myeloid progenitor (CMP), megakaryocyte-erythroid progenitor (MEP), dendritic cell (DC).

HSPC Stem/Multi, LMPP, and CLP remained persistently depleted from PreTx and EI through post-transplant follow-up (**Fig. 4A; Supp. Fig. 4A**), indicating sustained constraint of upstream stem and lymphoid-entry capacity. Within these primitive compartments, interferon and ISG signature pathways and genes were highest at PreTx, decreased by EI, then re-emerged post-transplant, followed by more lineage-restricted behavior at later timepoints (**Fig. 4B**; **Fig. 4F; Supp. Fig. 4F; Supp. Table 6**). In parallel, AP-1/IEG programs were broadly reduced at PreTx, including *FOS*/*FOSB*, *JUN*/*JUNB*/*JUND*, and *DUSP1*/*DUSP2*, with modest changes through EI and significant recovery across multiple progenitor subsets by ASCT1y (**Fig. 4D-E**; **Supp. Fig. 4B, 4E**). Dissociation and handling artifacts were assessed and were inconsistent with this pattern (**Supp. Fig. 4G**), supporting a disease-associated rather than technical origin. Together, these findings suggest that persistently depleted upstream progenitors occupy a transcriptionally distinct state during disease and early recovery.

CMP Core and DC progenitor subsets transiently rebounded at EI or ASCT90d, followed by later decline (**Fig. 4A; Supp. Fig. 4A**). ISG signatures in this trajectory were higher at PreTx than in the persistently depleted subsets, decreased to normal levels then higher than normal post-transplant with lineage-specific variability (**Fig. 4F; Supp. Fig. 4F; Supp. Table 6**). Within the DC axis, cDC and pDC progenitor subsets diverged from EI to ASCT1y, including state-specific behavior in IRF3 pathway signatures that was most evident in Prog DC pDC (**Fig. 4B**), consistent with non-uniform recovery within antigen-presenting lineages despite shared niche-level pressures.

Committed B-lineage progenitors expanded progressively during recovery (**Fig. 4A; Supp. Fig. 4A; Supp. Table 6**). This expansion occurred alongside heterogeneous metabolic and proliferative remodeling. OXPHOS was elevated or near healthy at PreTx and EI in multiple progenitor states, then became broadly suppressed post-transplant, with relative preservation in erythroid-biased progenitors (**Fig. 4B**). In contrast, FOXM1 and other cell-cycle pathways were strongly lineage-restricted and most consistently induced in proliferative B-lineage progenitors, erythroid states and proliferative monocytes (**Fig. 4B; Supp. Fig. 4C**), and these energetic and proliferative modules were frequently uncoupled within specific states (**Fig. 4B**). This uncoupling extended to DNA repair and cell-cycle coordination where early progenitors maintained high DNA repair signatures with heterogeneous cell-cycle activity. After transplant DNA repair signatures shifted toward healthy levels while many states remained compositionally depleted, indicating that depletion is not explained by a single uniform DNA repair-proliferation signature (**Supp. Fig. 4D**; **Fig. 4A**). Finally, BMIF proteomic profiling of canonical niche growth factors (CSF3, IL11, KITLG, THPO) showed substantial inter-patient variability without a consistent cohort-wide trajectory (**Fig. 4G**).

### Myeloma, therapy and transplant compartmentalize cytotoxic T and NK remodeling between marrow and blood

Cytotoxic remodeling emerged as a shared feature of disease and therapy in both marrow and blood (**Fig. 2H, 2J**). To compare compartments directly, we aligned cytotoxic CD8 and NK states using canonical effector modules (**Fig. 5A**). CD8 effector-memory states were resolved into two conserved modules, a *GZMK*+ *GZMH*+ *CD27*+ module and a *GZMB*+ *PRF1*+ *CD27*-module, and NK subset labels aligned across compartments when present (**Fig. 5A**). Despite this conserved structure, trajectories differed by compartment. In marrow, effector states remained enriched and recovery of naïve CD8 T cells was incomplete through ASCT2y. In blood, changes were dominated by a post-transplant expansion of multiple CD8 effector-memory states with partial normalization by ASCT2y, and durable reduction of naïve CD8 T and GZMK-EM post-transplant (**Fig. 5B-C; Supp. Table 5,6).** NK subset dynamics were more pronounced in the blood than in the marrow, with treatment-related expansion of CD56br and CD56dim GZMK+ NK cells, while marrow populations were stable **(Supp. Fig. 5A-B**). Pseudobulk differential expression analysis showed that cytotoxic machinery was consistently higher in marrow at PreTx and EI than in healthy controls, most prominently in CD8 T EM2 (*PRF1*, *GZMB*, *GZMH*) and in CD8 T EM1 and tissue-resident subsets with a *GZMK*-biased module, while matched PBMC effector-memory states were similar to, or lower than, healthy controls (**Fig. 5D**). These results indicate that blood captures the identity of cytotoxic states but underestimates the magnitude of effector activation at the disease site.

**Figure 5:**
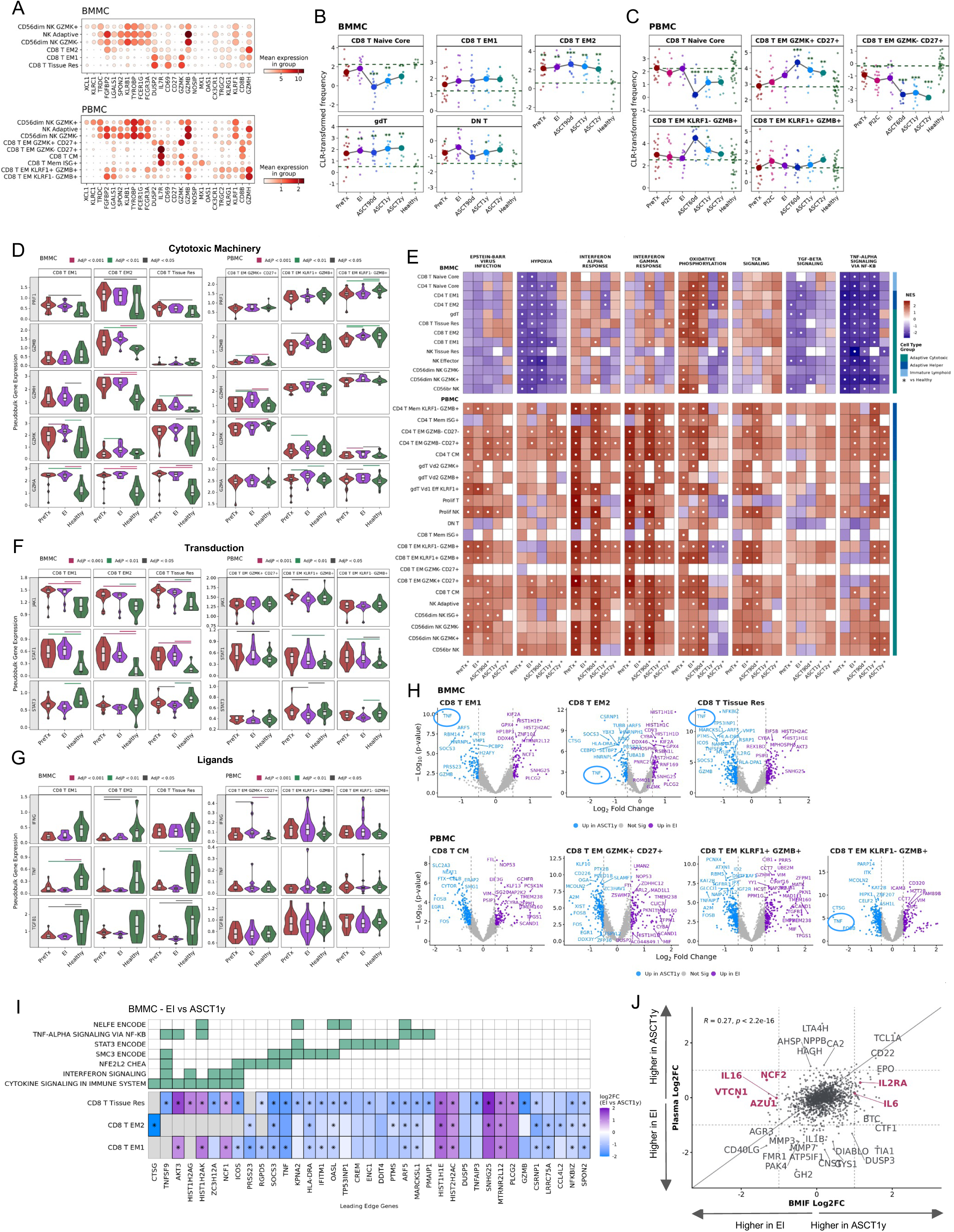
Cytotoxic T and NK state alignment, trajectories and pathways in marrow and blood. **(A)** Marker-gene dot plots defining aligned cytotoxic CD8 T and NK states in BMMCs (top) and PBMCs (bottom). **(B-C)** Longitudinal CLR-transformed abundances of cytotoxic T cell states in BMMCs **(B)** and PBMCs **(C)** across treatment timepoints relative to healthy donors. **(D)** Pseudobulk expression of cytotoxic machinery genes across cytotoxic CD8 subsets in BMMCs (left) and PBMCs (right). **(E)** FGSEA pathway enrichment (NES) across cytotoxic T and NK subsets at each timepoint relative to healthy controls, shown separately for BMMCs and PBMCs. White dots denote AdjP<0.05. **(F-G)** Pseudobulk expression of signaling and transduction genes **(F)** and effector ligands and trafficking markers **(G)** across cytotoxic CD8 subsets in BMMCs and PBMCs. **(H)** EI versus ASCT1y pseudobulk differential expression in cytotoxic CD8 subsets in BMMCs (top) and PBMCs (bottom). **(I)** Differentially expressed genes from EI versus ASCT1y comparisons showing pathway membership and gene-level log2FC across CD8 subsets. **(J)** Concordance of EI versus ASCT1y protein log2 fold changes between BMIF and plasma.

Consistent with this compartmental divergence in effector output, FGSEA revealed a parallel inversion in metabolic and inflammatory signatures (**Fig. 5E; Supp. Fig. 5F-G; Supp. Table 5,6**). Marrow cytotoxic states showed suppressed hypoxia-associated transcription and attenuated TNF-α signaling via NF-κB, whereas circulating cytotoxic subsets showed sustained expression of both signatures throughout therapy, indicating that inflammatory activation in the blood is not normalized by treatment and does not reflect cytotoxic state in the marrow niche. Interferon response signatures were similarly compartment specific, in marrow IFN programs were overall suppressed but showed treatment responsive remodeling, with ISG reductions and HLA class II upregulation by EI (**Supp. Fig. 5D**). Viral-response gene sets increased at EI in selected γδ T and CD8 effector-memory and tissue-resident states (**Fig. 5E**, **Supp. Fig. 5D**). In contrast, blood cytotoxic subsets maintained elevated IFN signatures across all timepoints, with leading edge genes dominated by HLA genes including HLA class II rather than reductions in ISG genes (**Fig. 5E**, **Supp. Fig. 5E**). Marrow CD8 cytotoxic states had higher *JAK1* and *STAT1* and lower *STAT3* than healthy controls, whereas PBMC differences were smaller and subset-dependent (**Fig. 5F**)^41^. Together, these results indicate that marrow cytotoxic IFN and inflammatory programs are disease-suppressed and treatment-sensitive, whereas their blood counterparts are constitutively elevated and treatment-resistant.

To connect compartmental signatures to functional output and residency, we compared ligands, trafficking markers, and inhibitory and differentiation-associated transcripts in matched cytotoxic subsets. In marrow, CD8 effector-memory and tissue-resident states, *IFNG*, *TNF*, and *TGFB1* were reduced relative to healthy controls, whereas PBMC effector-memory states were closer to healthy (**Fig. 5G**). Marrow cytotoxic subsets showed a retention-associated profile, with higher *CCL5*, lower *CCR7*, and higher ITGAE relative to healthy (**Supp Fig. 5H**). Flow cytometry confirmed elevated CD69 protein on marrow CD8 EM and TEMRA subsets through EI, consistent with tissue retention despite variable CD69 transcript levels across scRNA-seq-defined subsets (**Supp. Fig. 5C**; **Supp. Fig. 5H**). In contrast, PBMC showed stable *CCL5* and elevated *CD69* without accompanying ITGAE or CCR7 changes, suggesting activation rather than a canonical tissue residency program (**Supp. Fig. 5I**). Consistent with a more inhibitory and terminally differentiated marrow phenotype^42^, *EOMES*, *TOX*, and *TIGIT* were higher in marrow through ASCT1y, whereas PBMC changes were more variable, smaller, and subset specific (**Supp. Fig. 5J-K**). Additional checkpoint and exhaustion-associated markers were similarly compartment-skewed, where in marrow subsets, *BTLA*, *CD160*, *ENTPD1, CD244, LAG3, and CD96* remain elevated relative to healthy, whereas *PDCD1* showed no significant difference (**Supp. Fig. 6A-B**), consistent with reports that MM does not uniformly induce PD-1-dominated terminal exhaustion^43^.

To localize recovery dynamics, we compared EI to ASCT1y at the gene and protein level (**Fig. 5H-J**). In marrow cytotoxic CD8 states, EI versus ASCT1y expression changes suggested partial reactivation, including increased *TNF* and *ICOS* alongside induction of NF-κB feedback regulators *SOCS3*, *NFKBIZ*, and *TNFAIP3* (**Fig. 5H-I; Supp. Fig. 5L**)^44,45^. PBMC changes were more subset-restricted, although the *KLRF1*-*GZMB*+ EM state showed increased *TNF* at ASCT1y, aligning most closely with marrow EM2 (**Fig. 5H**). In parallel, BMIF proteomics shifted from an induction-associated myeloid and immunoregulatory profile at EI, including AZU1, NCF2, VTCN1, and IL16, toward elevated cytokines and chemokines linked to T cell activation and recruitment by ASCT1y, including IL2RA and IL6 (**Fig. 5J; Supp. Fig. 5M**)^46,47^. These gene and protein changes are consistent with gradual niche remodeling from EI to ASCT1y, with partial reactivation of marrow cytotoxic subsets and a more T cell-permissive cytokine and chemokine environment.

### Myeloma and Treatment Effects on B cells

Malignant MM plasma cells secrete or induce cytokines, chemokines, and growth factors that promote their growth and survival, and have effects on non-malignant B cells. At presentation, the frequencies of most non-malignant B cell subsets in peripheral blood were modestly decreased compared to healthy controls (**Fig. 6A**; **Supp. Table 5**). VRd induction further decreased many of these (**Fig.6A**), and spectral flow cytometry confirmed these findings (**Fig. 6B**; **Supp. Fig. 3A**; **Supp. Table 3A**). Transitional and memory B cell subsets were among those most affected by VRd treatment. In contrast, non-malignant plasma cells remained near healthy levels^48^. After HDM and ASCT, B cell reconstitution occurred by normal ontology. Transitional B cell frequencies rebound above normal by 60 days post-ASCT and normalize by ASCT1y while naïve B cells were initially low but approached healthy levels by ASCT1y (**Fig. 6A-B**; **Supp. Table 5**). Memory subsets remained markedly reduced through ASCT2y, despite expectations of post-ASCT memory B cell recovery (**Fig. 6A**; **Supp. Table 5**)^49,50^. Because maintenance therapy resumed after transplant (**Supp. Table 1E**), it was impossible to distinguish disease-intrinsic, from treatment-related contributions to delayed memory recovery.

**Figure 6:**
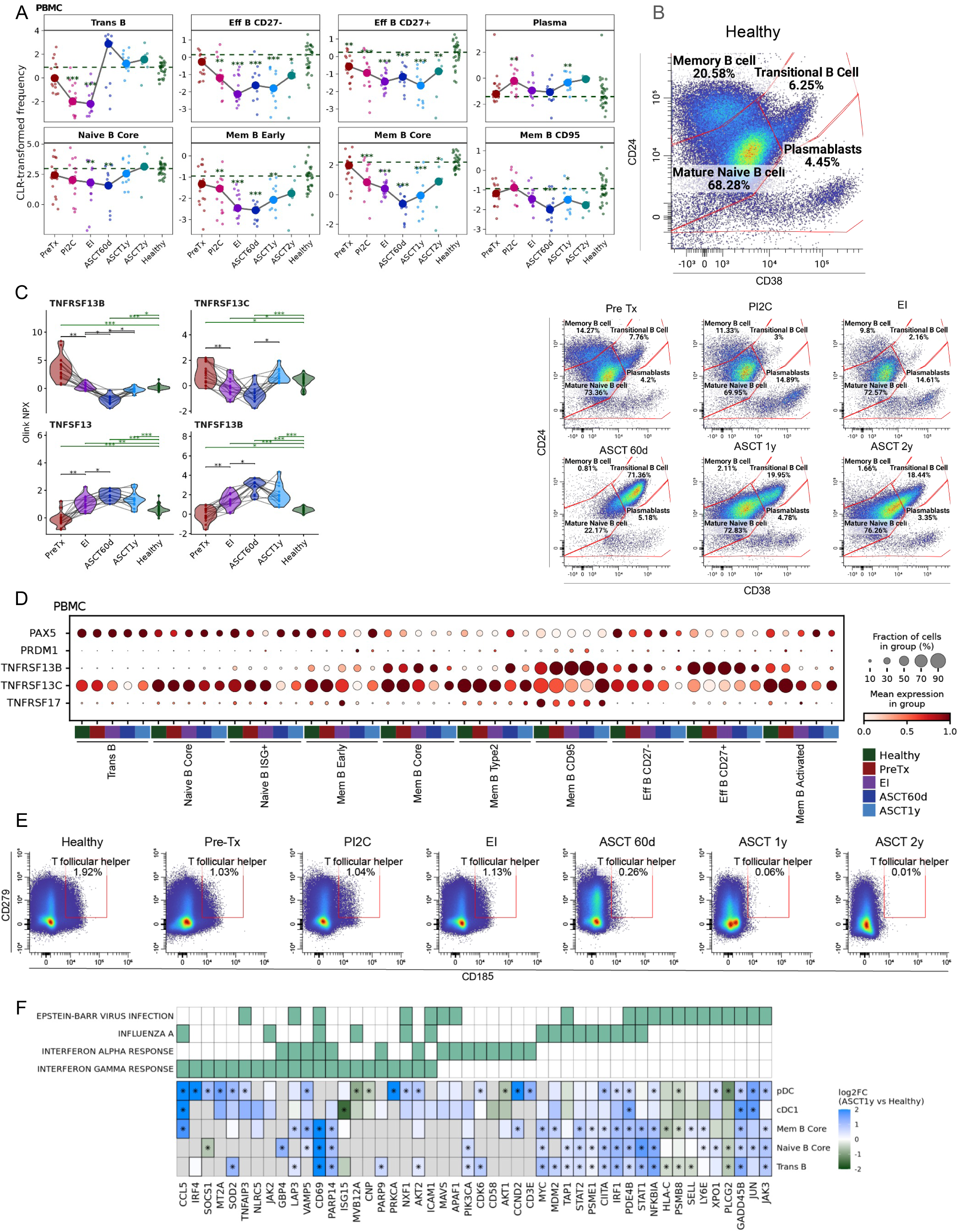
B cell subset frequencies, growth factors and Tfh context across therapy. **(A)** CLR-transformed frequencies of PBMC B cell subsets across timepoints relative to healthy controls. **(B)** Concatenated PBMC flow cytometry gating of B cell maturation states across induction and post-ASCT recovery, with healthy donors shown for reference. **(C)** Plasma proteomics of BAFF/APRIL axis components across timepoints. Black lines denote longitudinal comparisons, green lines denote comparisons with healthy samples. **(D)** Expression of B cell differentiation markers and BAFF/APRIL receptors across PBMC B cell subsets and timepoints. **(E)** Concatenated flow cytometry gating of circulating T follicular helper cells across timepoints. **(F)** Pathway leading-edge genes differentially expressed in PBMC subsets at ASCT1y versus healthy donors.

B cell growth and maturation could be influenced by dysregulated expression of the trophic factors APRIL (TNFSF13) and BAFF (TNFSF13B). Plasma proteomics showed that circulating APRIL and BAFF were reduced at presentation (PreTx) relative to healthy controls, then increased during induction therapy and ASCT as malignant and non-malignant B cells were depleted, likely causing B cell lymphopenia and decreased ligand consumption (**Fig. 6C**)^51,52^. These levels declined as B cell numbers recovered post-ASCT but remained elevated beyond ASCT1y, consistent with incomplete memory B cell reconstitution (**Fig. 3A-B; Supp. 3A; Fig. 6A,C**) and a potential risk for promoting tumor recurrence. Despite these changes in circulating BAFF protein, BAFF transcript is highest in ISG+ myeloid subsets including CD14+ monocytes and cDC2’s in peripheral blood **(Supp. Fig. 7G**) and is relatively stable across therapy (**Supp Fig. 7G**; **Fig. 6C**). APRIL protein followed a similar pattern as BAFF in peripheral blood (**Fig. 6C**), but APRIL transcript was negligible across circulating immune subsets suggesting a non-hematopoietic or tissue source (**Supp. Fig. 7G**). BAFF and APRIL receptor subunit transcript expression was B cell maturation-stage specific at presentation, similar to that of healthy controls. CD95+ memory B cells and to a lesser extent, core memory B cells show levels similar to healthy controls at diagnosis, but varied with therapy (**Fig. 6D**). BAFF-R (*TNFRSF13C*) decreased at EI and ASCT60d with partial recovery by ASCT1y, TACI (*TNFRSF13B*) is enriched in antigen-experienced B cells and transiently increased during therapy, and BCMA (*TNFRSF17*) was restricted to mature CD95+ memory cells and increased through therapy (**Fig. 6D**).

**Figure 7:**
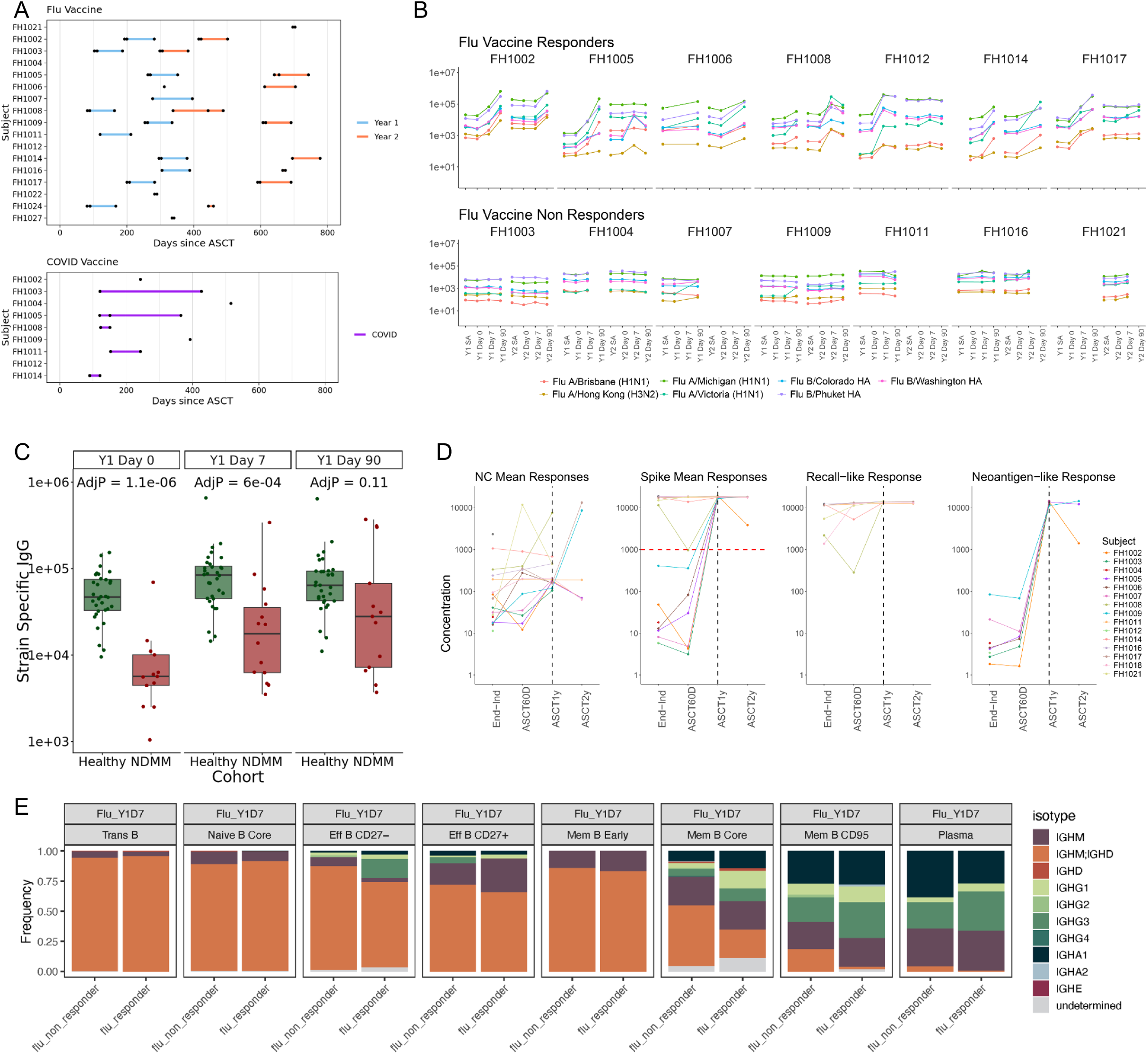
Vaccines and Functional Reconstitution. **(A)** Timing of seasonal influenza vaccination (two seasons) and COVID vaccination relative to ASCT for each subject. **(B)** Total influenza serology across baseline and post-vaccination sampling in Year 1 and Year 2, shown per subject and stratified by flu vaccine responder status (colored lines denote antigen-specific responses; timepoints as labeled). **(C)** Strain-specific IgG responses to influenza B/Phuket hemagglutinin measured by MSD at Day 0, Day 7, and Day 90 post-vaccination in NDMM patients versus healthy donors. Bonferroni-corrected p-values are shown for each timepoint. **(D)** SARS-CoV-2 vaccine serology across treatment timepoints in NDMM subjects, grouped by antigen class with lines connecting longitudinal samples. **(E)** Immunoglobulin isotype composition within PBMC B cell subsets at Day 7 post flu vaccination,

T cell help by CD40 ligand (CD40L) expressing T cells, particularly Tfh cells, is required for effective B cell class-switching and establishment of long-lived class-switched memory B cells^53^. Most T cell subsets decrease in frequency during induction therapy but post-ASCT, there is a strong drive to recover T cell numbers so there is a general skewing of T cells away from naive to more mature subsets by 2 years post-transplant (**Supp. Fig. 3A**). Tfh cells are a notable exception, remaining stable through induction therapy but profoundly depleted by myeloablative conditioning, and virtually absent through ASCT2y (**Fig. 6E**). Antigen presenting cell populations are unlikely to account for these differences, as peripheral DC subset frequencies are at or near healthy control levels throughout (**Supp. Fig. 7F**).

One year post-ASCT (ASCT1y), non-malignant B cell subsets from MM patients remain transcriptionally perturbed compared to B cells from healthy individuals. GSEA analysis demonstrated significant enrichment of Interferon Alpha and Gamma Response pathways in memory B core, naive B core, and transitional B cells, with Epstein-Barr Virus Infection (EBV) showing upregulation but not reaching significance (**Fig. 3D**). Leading edge genes in these pathways showed that by ASCT1y, B cell subsets retained an immune-primed program marked by increased expression of IFN responsive genes involved in antigen presentation and MHC processing (*CIITA, TAP1, PSME1, IRF1*), JAK/STAT signaling (*STAT1, STAT2, JAK3*) and cycling/stress response (*MYC, MDM2, GADD45B, JUN, PARP14*; **Fig. 6F; Supp. Table 5B**). Decreased expression of *PLCG2* in transitional and naive subsets post-ASCT, relative to healthy controls, suggests that BCR signalling is dampened prior to antigen exposure^54,55^ and decreased HLA-C and PSMB8 expression in mature B cell subsets suggests a decreased ability to process and present antigen (**Fig. 6F**)^56,57^. Together, these data suggest that ongoing inflammatory signalling reprograms recovering, non-malignant B cells in MM to have decreased BCR signaling and decreased ability to process and present MHC class-I antigen even 1 year post-ASCT^58,59^, adding to the evidence for ongoing B cell dysfunction after transplant.

### SARS-CoV-2 mRNA vaccination elicits robust responses while influenza responses remain variable after ASCT

Based on the lack of memory B cell and Tfh recovery, we hypothesize that patients may be left with an immunoglobulin class-switch defect for at least 2 years post-ASCT. To functionally test this, we evaluated immune responses to two vaccines administered as standard of care following ASCT, including 1) non-adjuvanted, high-dose influenza vaccine that contains a higher dose of flu antigen than standard flu vaccines^60,61^ and 2) COVID lipid nanoparticle mRNA vaccine in which the nanoparticles have been shown to act as a potent adjuvant that enhances immune responses (**Fig. 1A**; **Fig. 7A**)^62–64^.

We evaluated strain-specific IgG after post-ASCT high-dose influenza vaccination and found heterogeneous responses. Half of patients (7/14) mounted a multi-strain response in at least one year, whereas the remainder showed minimal responses (**Fig. 7B**). Overall IgG magnitude was reduced relative to age and sex matched healthy donors, illustrated by lower anti-B/Phuket IgG that has been in seasonal influenza vaccines from the 2015-16 to 2023-24 flu seasons (**Fig. 7C**). Flu vaccine responsiveness was not associated with sex, age, flu season, or vaccination timing relative to ASCT (**Supp. Fig. 7A,C-E**). Since the assay for evaluating strain-specific flu responses measures only IgG, we evaluated functional immunoglobulin class switching. We compared productive immunoglobulin transcripts at Day 7 and found that non-responders were less likely to switch from IgM or IgM/IgD to IgG isotypes, most evident in CD27-effector core memory, and CD95+ memory B cell populations (**Fig. 7E**).

Enrollment for this study began before emergency use authorization for the first COVID-19 mRNA lipid nanoparticle vaccines. This allowed assessment of neoantigen and recall responses after ASCT. Early participants were first vaccinated post-transplant, whereas later participants were vaccinated before NDMM diagnosis and again after ASCT (**Fig. 7A**). Because many doses were administered at community sites, complete dose histories were unavailable. Regardless of timing, all patients showed IgG responses to all tested spike receptor binding domains by ASCT1y (**Fig. 7D**; **Supp. Fig. 7B**), and four patients had nucleocapsid reactivity consistent with prior infection. Together, these results suggest that post-ASCT constraints can limit responses to non-adjuvanted influenza vaccination in some patients, whereas mRNA and lipid nanoparticle platforms provide sufficient innate stimulation to overcome these constraints.

## Discussion

Recent profiling of long-term survivors emphasized that bone marrow immune alterations can persist for years following therapy, supporting the concept of durable marrow changes even in the absence of residual disease^6^. That work established durability, but it does not define when these alterations emerge during therapy, or whether they are visible in blood. Our longitudinal design across PreTx, EI, and post-ASCT recovery resolves this, and three conclusions emerge. **First,** persistence of tumor cells after therapy is associated with malignant transcriptional states not fully captured by cytogenetics alone, consistent with selection of residual tumor states. **Second**, immune remodeling is compartmentalized, with marrow and blood showing opposing inflammatory and metabolic signatures despite overlapping cell identities, reflecting both compositional differences and compartment-specific regulation of cell states. **Third**, persistent immune dysfunction extends beyond transient cytopenia and points to constrained recovery affecting hematopoietic output, antigen presentation, and humoral memory maturation with downstream consequences for vaccine responsiveness. These constraints are reflected in lineage-biased progenitor trajectories and durable cytotoxic skew in the marrow.

Clinical risk stratification in MM relies heavily on cytogenetics, but therapy acts on functional states that can span genotypes. Our data support transcriptional state as an additional organizing facet because several shared malignant states contract with induction, while others persist across patients regardless of tumor cytogenetic background. The key conceptual point is that a state that has low abundance at baseline but becomes enriched after treatment is being selected for by therapy. In our cohort, the most striking example of this is NDMM_5, which becomes enriched among residual tumor cells across individuals despite low abundance at diagnosis, providing a reference point for thinking about composition of residual disease after induction. This state-based view extends residual disease interpretation beyond quantity alone. Even when tumor burden is low by standard measures, the residual states can differ from pre-treatment and may impose a different set of pressures on the marrow immune ecosystem during recovery. In our cohort, features of more persistent shared states were consistent with stress tolerance and metabolic flexibility, supporting a model where induction debulks therapy-sensitive states while enriching residual states that may require sequential or combination strategies tailored to state heterogeneity and plasticity. Persistence of transcriptionally defined tumor states across cytogenetic backgrounds suggests that MRD monitoring and therapeutic strategies could be strengthened by incorporating cell state profiling alongside standard cytogenetic testing. We also observed a reciprocal relationship between MYC-associated signatures and the SUZ12 ENCODE signature, raising the possibility that malignant plasma cells occupy alternative chromatin configurations with distinct transcriptional constraints. Because SUZ12 is a core component of PRC2, these opposing pathways may reflect differences in chromatin repression and transcriptional output. This axis provides a foundation for follow up studies that ask whether different residual states differ in sensitivity to therapies that modulate transcriptional output or chromatin repression^19^.

A defining feature of this study is that marrow and blood are not interchangeable windows into immune status at diagnosis and through therapy. Even when the same broad cell subsets are present, marrow and blood occupy different and often opposing transcriptional states. This supports the idea that the marrow niche imposes local constraints on the immune state, with caution that blood based monitoring can misrepresent disease site immune biology during recovery. The most consistent example is the compartment-wide inversion of metabolic and inflammatory signatures, which is consistent across most immune cell subsets and also includes most malignant plasma cell states. In healthy individuals, marrow is a relatively low-oxygen tissue, with local gradients and well described metabolic adaptations^65^. In MM marrow, hypoxia sensing transcriptional signatures and inflammatory gene sets were broadly suppressed across progenitor and immune subsets, whereas peripheral blood showed a distinct landscape dominated by inflammatory signaling across therapy relative to healthy. This inversion is not readily explained by systemic cytokine levels alone, and instead points to tissue-specific regulation imposed by the myeloma marrow environment and its remodeling during therapy.

These compartmental differences have consequences for biomarker strategy and therapeutic targeting. Blood and marrow report different aspects of the immune system during treatment, and blood-based correlates may not capture marrow immune suppression. This motivates interpreting biomarkers by compartment and modality, with paired sampling used to distinguish shared systemic signals from marrow-restricted effects. EI captures a low tumor burden, therapy-exposed state, whereas ASCT1y reflects hematopoietic regeneration and a recovering microenvironment. For example, VTCN1 was enriched in BMIF at EI, placing this candidate immune checkpoint ligand in the marrow and motivating prospective evaluation as an immunotherapeutic target in MM.

A key mechanistic question is whether persistent abnormalities reflect altered hematopoietic output, depletion of key lineages, durable impairment of cell state, or some combination. Our data indicate contributions from all three. Persistent post-therapy abnormalities reflect both incomplete lineage recovery and durable functional state shifts in reconstituting cells, with only partial concordance between numerical recovery and transcriptional state. This is evident in the lineage biased progenitor trajectories, durable expansion of cytotoxic subsets, and divergence between compositional and transcriptional recovery across lineages. The three progenitor trajectories indicate constrained upstream capacity with selective expansion of committed lineages during recovery, consistent with evidence that early HSPC and lymphoid progenitor constraints can shape downstream immune composition, even when compartments partially normalize^66,67^. Early HSPC and early lymphoid states remain persistently depleted and transcriptionally abnormal, with blunted AP-1/IEG signatures and sustained attenuation of hypoxia-associated signatures. This is notable given evidence that hypoxia-associated pathways support HSC function and marrow niche fitness^68,69^, and raises the question of whether the post-therapy marrow niche supports the same quality of hematopoietic output as before disease. Given that consolidation and transplant follow soon after induction, the status of hematopoietic stem and progenitor lineages at EI, which influences mobilization and graft composition, is likely to influence post-ASCT immune reconstitution^70,71^. Coordinated attenuation of inflammatory signaling across progenitor subsets, including reduced TNFα via NF-κB together with delayed recovery of AP-1/IEG signatures, suggests that recovery proceeds with broad suppression of pathways normally engaged by cellular and microenvironmental stress. The basis of the hypoxia signature remains to be defined and motivates targeted mechanistic studies to determine whether hypoxia signature suppression reflects differences in niche oxygen tension, altered transcriptional responsiveness, or shifts in state composition.

Cytotoxic remodeling provides an independent readout of marrow microenvironmental influence. Persistent skewing toward effector states in the marrow, together with differences in cytotoxic, inhibitory, and differentiation associated features aligns with prior reports that myeloma promotes dysfunction across T and NK subsets^72–74^, while not uniformly inducing classical terminal exhaustion states^43^. We further find that the suppression of hypoxic, inflammatory and stress-response signatures extends from hematopoietic subsets to cytotoxic subsets in the marrow compartment, whereas corresponding circulating cytotoxic subsets show increased pathway signatures. Relative to blood, marrow cytotoxic states have reduced *IFNG*, *TNF*, and *TGFB1*, together with increased inhibitory and exhaustion-associated markers. The gradual changes in cytotoxic gene transcription between EI and ASCT1y is consistent with the slow reconstitution of the marrow immune microenvironment after transplant that does not fully return to healthy levels within the follow-up window^49,75^. Even without uniform induction of all classical canonical terminal exhaustion markers, marrow cytotoxic EM subsets show a constrained activation state, with reduced engagement of inflammatory and stress-response signatures, consistent with a niche-imposed attenuated EM state.

The marrow constraints described above plausibly contribute to compositional and functional defects observed in the periphery across therapy. The immune landscape at EI is both cytopenic and functionally altered, marked by loss of naïve and memory B subsets and Tfh subsets together with the expansion of ISG+ and IL1B+ inflammatory monocytes in the blood. IL1B+ monocytes remained elevated through post-ASCT recovery, and BAFF and APRIL protein levels were high in blood and marrow, consistent with persistent inflammatory and trophic signals that can bias lymphoid reconstitution away from mature memory B and Tfh recovery. In addition to compositional defects, MM patients who fail to respond to influenza vaccination show more prominent unswitched IgM and IgM/IgD across memory B cell subsets at day 7 post-vaccination, whereas responders show higher IgG and IgA isotypes, indicating a more mature switched memory B cell population in responders at the time of vaccination. Together these findings point to a higher threshold for germinal center-dependent antibody responses in the setting of weak innate stimulation.

This threshold effect is evident in the vaccine data. Vaccination is a standard component of post-transplant care, but immunogenicity varies substantially across vaccine platforms in MM (Fred Hutch LTFU Guidelines, Table IX.A2)^76^. In our cohort, influenza responses were heterogeneous with stable responder and non-responder patterns across seasons, whereas SARS-CoV-2 mRNA vaccination elicited robust serologic responses. The influenza vaccine used was high-dose but not adjuvanted whereas the SARS-CoV-2 mRNA vaccination is delivered in lipid nanoparticles (LNP) that provide adjuvant-like innate stimulation^64^. We interpret this contrast as evidence that persistent post-ASCT impairment in B and T cell function required for germinal center responses creates a higher threshold for achieving functional neutralizing antibody responses, which may be met by mRNA/LNP platforms and not by non-adjuvanted influenza vaccine in a subset of patients. The adjuvant dependence of vaccine success provides direct rationale for clinical trials comparing adjuvanted (or mRNA based influenza platforms) versus high-dose non-adjuvanted influenza vaccine in post-transplant MM patients.

### Study limitations and conclusions

Although the cohort size is modest and later timepoints include attrition, baseline risk distribution and responses are consistent with published NDMM VRd with ASCT cohorts. Evolving standard of care therapy with increasing use of daratumumab based regimens (DVRd) limited power for subgroup analyses and clinical associations. Pathway analyses reflect transcriptional signatures rather than direct functional measurements, for example, *HIF1α* expression and suppressed hypoxia pathway signatures cannot distinguish true differences in niche oxygen tension from upstream transcriptional silencing. Future MM related studies will be important to directly test niche mechanisms that sustain marrow-local immune remodeling. Finally, influenza comparisons to external healthy controls are complicated by vaccine formulation differences and variable timing post-transplant. Despite these constraints, paired compartment sampling and orthogonal modalities strengthen inference about compartment-specific signatures and therapy related remodeling.

Our findings support a model in which myeloma and its treatment establish a prolonged niche-constrained state of immune reconstitution. Therapy selects among shared malignant plasma cell states beyond what is captured by cytogenetics and leaves marrow and blood at divergent inflammatory and metabolic baselines that persist throughout recovery. Lineage-biased progenitor trajectories and sustained transcriptional abnormalities in reconstituting immune cells indicate that this persistent dysfunction is not explained by depletion alone, but reflects constrained hematopoietic output and slow normalization of marrow immune function. Whether this constrained marrow function is due to MM or treatment “scarring” or due at least in part to ongoing maintenance therapy could not be distinguished in this study due to the use of maintenance therapy as a standard of care for post-ASCT patients. Variable post-transplant influenza responses alongside robust SARS-CoV-2 mRNA vaccine responses suggest that the vaccine platform can overcome some disease and therapy related immune constraints and point toward adjuvanted strategies as a practical intervention to improve infection-related morbidity in MM.

## Supporting information

Supplemental Table 7_reagent table

## Resource availability

### Data availability

Single-cell RNA-seq data have been deposited in GEO under accession numbers **GSE309595** (PBMC scRNA-seq) and **GSE309593** (BMMC CITE-seq). Primary analyses in this manuscript focus on the VRd cohort, with exploratory analyses of DVRd as described in the Results. The deposited dataset includes samples from the primary VRd cohort and additional samples from the exploratory daratumumab plus VRd (DVRd) cohort and a relapsed refractory multiple myeloma (RRMM) cohort, which are provided as part of the resource and data release. Clinical metadata for all deposited samples are provided in Supplemental Table S1. Raw sequencing data (FASTQ files) will be deposited in dbGaP and will be publicly available upon publication (accession pending). This study also analyzed publicly available datasets from GEO (**GSE253355**, **GSE271896**).

### Code availability

All code used to generate the analyses is available on GitHub and will be archived on Zenodo upon publication (links to be provided in the final version).

Link: https://github.com/aifimmunology/multimodal-myeloma

### Additional resources

Processed data (Flow, Olink, MSD), analysis objects, and clinical metadata are available through the Human Immune System Explorer (HISE) portal (link to be provided). Selected results can also be explored via an interactive viewer (https://apps.allenimmunology.org/aifi/insights/ndmm/). Additional information required to reanalyze the data reported in this paper is available from the lead contact upon request.

## Acknowledgments

We thank the study participants and the clinical team at Fred Hutchinson Cancer Center for dedicated support throughout the cohort sample collection. We thank all members of the Allen Institute for Immunology, specifically Leila Shiraiwa, Nina Estep, Nina Kondza and the facilities and operations teams for sample handling and research support; Stark Pister, Jessica Lang, and the Human Immune System Explorer (HISE) software development team for data related support. The authors thank the Allen Institute founder, Paul G. Allen, for his vision, encouragement, and support. Research reported in this publication was supported by the Allen Institute. Overview figures were created with BioRender.com

## Author Contributions

**Conceptualization:** S.R., T.F.B., E.W.N., P.D.G., L.T.G., D.J.G., T.R.T. and M.L.A.-H. **Data Curation:** M.A.D., M.S.K., X.S., U.K., L.Y.O., L.T.G. and M.L.A.-H. **Formal Analysis:** A.C., S.R.Z., M.A.D., P.C.G., N.M., P.I.M., K.J.L., J.R., M. Singh, Y.D.H., L.Y.O., T.P., C.G.P., Z.J.T. and L.T. **Funding Acquisition:** T.F.B., S.M.H., A.W.G. **Investigation:** M.A.D., P.C.G., K.J.L., J.R., V.H., J.G., E.W.D., A.T.H., E.K.K., C.L., R.R.M., B.M., V.P., C.G.P., C.R.R., E.G.S., T.J.S., M.D.A.W., P.J.W. and S.D.A.-S. **Methodology:** A.C., S.R.Z., M.A.D., P.C.G., N.M., E.L.K., M.C.G., D.G., Y.D.H., P.D.G., X.L. and L.T. **Project Administration:** A.C., S.R.Z., M.S.K., X.S., L.A.B., L.T.G., M. S., T.R.T. and M.L.A.-H. **Resources:** P.M., M.K. and D.J.G. **Software:** C.M.L., G.L.K., P.M. and L.T.G. **Supervision:** S.R.Z., M.S.K., J.R., J.G., Y.D.H., P.M., Z.J.T., S.D.A.-S., E.M.C., L.A.B., T.F.B., A.W.G., E.W.N., X.L., S.M.K., P.J.S., M. S., M.K., D.J.G., T.R.T. and M.L.A.-H. **Validation:** P.C.G. **Visualization:** A.C., S.R.Z., M.A.D., N.M. and M.L.A.-H. **Writing-Original Draft:** A.C., S.R.Z., M.A.D., P.C.G., M.S.K., K.J.L., J.G., T.R.T. and M.L.A.-H. **Writing-Review & Editing:** E.L.K., X.L., S.M.K., L.T.G., L.T., T.R.T. and M.L.A.-H.

## Supplemental Information

Document S1. Figures S1-S7

Supplemental Tables S1-S6 contain cohort metadata, cytogenetic data, flow cytometry results, Olink proteomics results, PBMC and BMMC pseudobulk RNA-seq results, and serology results.

Supplemental_Table_1_metadata.xlsx Supplemental_Table_2_cytogenetic_data.xlsx Supplemental_Table_3_FLOW.xlsx Supplemental_Table_4_OLINK .xlsx Supplemental_Table_5_RNAseq_PBMC_all_results.xlsx Supplemental_Table_6_RNAseq_BMMC_all_results.xlsx

## Supplemental Table Legends

**Supplemental Table 1: Patient Metadata.** This workbook contains clinical, demographic, and treatment metadata for all study participants. Sheet **A_FH1_Cohort** provides an overview and clinical data for all enrolled VRd and DVRd NDMM participants. Sheet **B_Healthy_PBMC_Cohort** describes the healthy peripheral blood donor cohort used as reference controls for PBMC analyses. Sheet **C_Healthy_BM_Cohort** describes the healthy bone marrow donor cohort used as reference controls for BMMC analyses. Sheet **D_FH1_Therapy** details the induction treatment regimens and dose information for VRd and DVRd participants. Sheet **E_FH1_MaintenanceTherapy** summarizes post-ASCT maintenance therapy details. Sheet **F_FH1_ORIG_Imaging** contains clinical imaging data including PET/CT, MRI, and skeletal survey findings. Sheet **G_FH2_Cohort** provides an overview of enrolled participants in the exploratory RRMM cohort. Sheet **H_FH2_Therapy** details the treatment regimens for DVRd cohort participants. Sheet **I_FH2_Risk_Response** summarizes disease risk stratification and clinician-recorded therapeutic response assessments for DVRd participants. Sheet **J_HRMM** contains high-risk multiple myeloma classifications using updated IMWG_HRMM criteria. Sheet **K_Tumor_Burden** provides longitudinal tumor burden measurements across treatment timepoints. Sheet **L_Demographics** summarizes demographic characteristics of the full study cohort. Sheet **M_NDMM_RRMM_KIR** contains killer cell immunoglobulin-like receptor (KIR) genotyping data for NDMM and RRMM participants. Sheet **N_NDMM_RRMM_HLA** contains human leukocyte antigen (HLA) genotyping data for NDMM and RRMM participants. Sheet **O_NDMM_RRMM_FCGR** contains Fc gamma receptor (FcGR) allele genotyping data for NDMM and RRMM participants.

**Supplemental Table 2: Cytogenetic Data.** This workbook contains cytogenetic profiling data for the NDMM cohort derived from clinical FISH and karyotype analyses. Sheet **A_cyto_data_cleaned** provides the cleaned and processed cytogenetic dataset for each subject used in downstream analyses. Sheet **B_filt_heatmap_data** contains the filtered cytogenetic dataset and mean expression for genes of interest for each subject used to generate the cytogenetic heatmap visualizations presented in the manuscript.

**Supplemental Table 3: Flow Cytometry.** This workbook contains spectral flow cytometry analysis from peripheral blood across treatment timepoints. **Sheet A_vrd_healthy_comparison** contains VRd versus healthy comparison results. **Sheet B_vrd_longitudinal_comparison** contains longitudinal comparison results for the VRd cohort. **Sheet C_dvrd_healthy_comparison** contains DVRd versus healthy comparison results.

**Supplemental Table 4:** Plasma and BMIF Olink Proteomics. This workbook contains pairwise differential protein expression analyses from longitudinal bone marrow and plasma samples. **Sheet A_boneMarrow_longitudinal_pairwise** contains pairwise differential protein expression results from longitudinal bone marrow comparisons. **Sheet B_plasma_longitudinal_pairwise_de** contains analogous pairwise differential protein expression results from longitudinal plasma comparisons.

**Supplemental Table 5: scRNA-seq PBMC — All Results.** This workbook contains single-cell RNA sequencing gene set enrichment and cell type compositional analysis results from peripheral blood mononuclear cells (PBMCs). Sheets prefixed fgsea contain Fast Gene Set Enrichment Analysis (FGSEA) results for the indicated pairwise comparison, including normalized enrichment scores (NES) and adjusted p-values across cell types and pathways. Sheets prefixed freq contain centered log-ratio (CLR)-transformed cell type frequency results for the indicated pairwise comparison, including effect sizes and statistical test results. Sheets are organized as follows: **!_PBMC_L3_label** catalogs all L3 label subsets used throughout this study. FGSEA sheets: **A_fgsea_ASCT1y_vs_ASCT2y**, **B_fgsea_ASCT1y_vs_Healthy**, **C_fgsea_ASCT2y_vs_Healthy**, **D_fgsea_ASCT60d_vs_ASCT1y**, **E_fgsea_ASCT60d_vs_Healthy**, **F_fgsea_EI_vs_ASCT1y**, **G_fgsea_EI_vs_ASCT2y**, **H_fgsea_EI_vs_ASCT60d**, **I_fgsea_EI_vs_Healthy**, **J_fgsea_PI2C_vs_EI**, **K_fgsea_PI2C_vs_Healthy**, **L_fgsea_PreTx_vs_EI**, **M_fgsea_PreTx_vs_Healthy**, **N_fgsea_PreTx_vs_PI2C**. Frequency/compositional sheets: **O_freq_all_timepoint_frequency** (CLR-transformed cell type frequencies across all timepoints), **P_freq_asct1y_vs_asct2y_results**, **Q_freq_asct1y_vs_healthy_result**, **R_freq_asct2y_vs_healthy_result**, **S_freq_asct60d_vs_asct1y_result**, **T_freq_asct60d_vs_healthy_result**, **U_freq_ei_vs_asct1y_results**, **V_freq_ei_vs_asct2y_results**, **W_freq_ei_vs_asct60d_results**, **X_freq_ei_vs_healthy_results**, **Y_freq_pi2c_vs_ei_results**, **Z_freq_pi2c_vs_healthy_results**, **AA_freq_pretx_vs_ei_results**, **AB_freq_pretx_vs_healthy_result**, **AC_freq_pretx_vs_pi2c_results**.

**Supplemental Table 6: scRNA-seq BMMC — All Results** This workbook contains single-cell RNA sequencing gene set enrichment and cell type compositional analysis results from bone marrow mononuclear cells (BMMCs). As in Supplemental Table 5, sheets prefixed fgsea contain FGSEA results for the indicated pairwise comparison and sheets prefixed freq contain CLR-transformed cell type compositional results. Additional specialized sheets are included for malignant plasma cell analyses. Sheets are organized as follows: **!_BMMC_L3_labels** catalogs all BMMC labeled subsets used throughout this study. Frequency/compositional sheets: **A_freq_all_timepoint_frequency** (CLR-transformed cell type frequencies across all BMMC timepoints), **B_freq_asct1y_vs_asct2y_results**, **C_freq_asct1y_vs_healthy_result**, **D_freq_asct2y_vs_healthy_result**, **E_freq_asct90d_vs_asct1y_result**, **F_freq_asct90d_vs_healthy_result**, **G_freq_ei_vs_asct1y_results**, **H_freq_ei_vs_asct2y_results**, **I_freq_ei_vs_asct90d_results**, **J_freq_ei_vs_healthy_results**, **K_fgsea_plasma_state_scores_summary** (pathway enrichment score summary across malignant plasma cell Leiden cluster states), **L_freq_pretx_vs_ei_results**, **M_freq_pretx_vs_healthy_results**. FGSEA sheets: **N_fgsea_ASCT1y_vs_ASCT2y, O_fgsea_ASCT1y_vs_Healthy**, **P_fgsea_ASCT2y_vs_Healthy**, **Q_fgsea_ASCT90d_vs_ASCT1y**, **R_fgsea_ASCT90d_vs_Healthy, S_fgsea_EI_vs_ASCT1y**, **T_fgsea_EI_vs_ASCT2y**, **U_fgsea_EI_vs_ASCT90d**, **V_fgsea_EI_vs_Healthy**, **W_fgsea_PreTx_vs_EI**, **X_fgsea_PreTx_vs_Healthy**. Plasma cell-specific sheet: **Y_FGSEA_Malignant_PC_to_Healthy_PC** contains FGSEA results directly comparing malignant plasma cells from NDMM participants to healthy donor plasma cells, identifying gene sets that are enriched or depleted in malignant versus non-malignant plasma cell states.

## STAR Methods

Reagent table: (Supp. Methods_ Reagent info.xlsx)

### Figure conventions and statistical reporting

#### Common figure conventions

Dot plots encode mean expression by color and the fraction of cells expressing the gene by dot size. Violin and box plots display sample-level or pseudobulk distributions as indicated in the plot titles. Heatmaps display either gene set enrichment scores (NES) or log2 fold changes, with direction as indicated by the color scale. Volcano plots show effect size on the x axis and multiple-testing-adjusted significance on the y axis, with thresholds as specified below. For asterisks and significance bars/lines, unless otherwise stated in the figure legends, adjusted P values are reported as AdjP; unless otherwise noted, significance is AdjP <0.05, with *, ** for AdjP <0.01 and *** for AdjP <0.001. For violin plots, horizontal lines indicate significance of the specified comparisons, and line colors denote AdjP thresholds as shown in the figure legend key: maroon (AdjP <0.001), seagreen (AdjP <0.01), gray (AdjP <0.05), unless otherwise noted.

### Study participants, treatment, and clinical assessments

A cohort of 25 (17 VRd; 8 DVRd) transplant-eligible patients aged 30 to 70 with newly diagnosed multiple myeloma (NDMM) requiring systemic therapy was enrolled in Seattle, WA (IRB #10265) with approval from the Institutional Review Board of Fred Hutchinson Cancer Center. Patients were followed prospectively through standard-of-care VRd induction, high-dose melphalan conditioning, autologous stem cell transplantation, consolidation, and maintenance therapy (Fig. 1A; **Table 1**). Induction therapy was administered for at least four cycles, followed by high-dose melphalan and ASCT, consolidation for two cycles, and lenalidomide maintenance thereafter; dose adjustments were made for medical indications (**Supp. Table 1D**). By institutional standard, a very good partial response was required to proceed to transplant, achieved through extension of induction, regimen modification, or both. Blood and bone marrow samples were collected at the following timepoints: prior to induction therapy (PreTx), after two cycles of induction therapy (PI2C, blood only), end of induction (EI), 60 days post-transplant (ASCT60d, blood only), and 90 days post-transplant (ASCT90d, bone marrow biopsy obtained between 60 and 100 days post-transplant at the clinical discretion of the treating physician) (Fig. 1A). Additional blood and bone marrow samples were obtained at one year (ASCT1y) and two years (ASCT2y) post-transplant based on sample availability (Fig. 1A). Diagnosis, treatment indication, and risk stratification were established using International Myeloma Working Group (IMWG) criteria with standard laboratory testing, pathology, and imaging, and disease activity was assessed each cycle (**Supp. Table 1**). Patients were staged using the revised international staging system (RISS) and were also retrospectively classified using updated IMWG high-risk criteria (IMWG_HRMM; Fig. 1B**; Supp. Table 1J**); high-risk status may be underestimated because TP53 mutation status was not available. ^77,78^.

This cohort was also followed for flu vaccine perturbation over the course of two consecutive flu years and blood samples were collected at day 0, day 7-9, and day 90+/-10 post-vaccination, as well as a baseline sample (a flu-series time point where the blood does not fall on the 0, 7, or 90 day timepoint).

### Clinical labs and imaging

A comprehensive clinical and laboratory profiling strategy was employed to capture both static and dynamic indicators of disease activity, immune status and therapeutic response. In addition to patient demographics, therapeutic response data (tumor burden and response), Whole body imaging with PET/CT to assess metabolic activity, complemented by MRI to evaluate bone marrow infiltration, and osseous survey or Xray for skeletal involvement, was used along with bone marrow evaluation to assess disease burden and monitor treatment response. Additionally, a set of laboratory measures was collected, obtained either at a single timepoint for those parameters expected to remain stable throughout therapy, or longitudinally at each visit to capture dynamic clinical changes (Supp. Table 1A). These included tumor burden, blood cell percentages and absolute counts, hematological indices, cytogenetic profiles, serum chemistries, cytomegalovirus (CMV) serostatus, serum protein electrophoresis (SPEP) and immunofixation electrophoresis (IFE), inflammatory markers such as erythrocyte sedimentation rate (ESR) and C-reactive protein (CRP), immunoglobulin levels, lipid profiles, and KIR, HLA, and FCGR allele status (Supp. Table 1A-B). Together, these clinical parameters provided a framework for evaluating patients heterogeneity and treatment-associated changes within and across our cohorts.

### Sample Collection

Fresh whole blood samples were collected from these subjects in heparinized syringes or sodium heparin vacutainers and processed to peripheral blood mononuclear cells (PBMCs) through a Ficoll-based approach. PBMCs were aliquoted in vials of 5×10^6^ cells/ml each and frozen in Cryo-storage media (90% FBS, 10% DMSO) within 4 hours of blood draw using a pre-chilled 4°C CoolCell or Mr. Frosty freezing container stored at -80°C for up to 72 hours before being transferred to liquid nitrogen for long-term storage.

Fresh whole blood samples were collected from NDMM subjects in K2 EDTA vacutainers or tubes and plasma supernatant samples were isolated using centrifugation, aliquoted, and frozen at -80°C within 4 hours of blood draw.

Bone marrow aspirate (BMA) samples were collected from NDMM subjects in heparinized syringes/sodium heparin vacutainers and bone marrow mononuclear cells (BMMCs) were isolated using a Ficoll-density gradient approach. Briefly, BMA samples were washed in equal volume with 1x PBS before layering the BMA/PBS mixture on top of an equal volume of Ficoll. BMMCs were aliquoted in vials each containing 1×10^6^ cells and frozen in Cryo-storage media (90% FBS, 10% DMSO) using a pre-chilled 4°C CoolCell or Mr. Frosty freezing container stored at -80°C for up to 72 hours before being transferred to liquid nitrogen for long-term storage.

Bone marrow interstitial fluid (BMIF) samples were collected from the supernatant of BMA samples, following centrifugation at 480xg for five minutes at room temperature. BMIF samples were aliquoted into 1.2 mL cryovials and stored at -80°C. Prior to sending samples to Olink for processing, samples were thawed on ice and aliquoted into 50 µL aliquots before being refrozen and stored at -80°C until shipment.

### Healthy Age Matched Donors

#### Sound Life Cohort (PBMC Samples)

Healthy adult donors aged 25-65 were prospectively recruited from Seattle, WA as part of the Sound Life Project, in collaboration with Benaroya Research Institute as previously described^23^ . Donors were excluded from enrollment if they had a history of chronic or autoimmune disease/infection or severe allergies. PBMC samples were collected from fresh whole blood using a Ficoll-based approach and frozen within 4 hours of blood draw. Plasma samples were processed, aliquoted and frozen within 4 hours of blood draw.

Matched healthy controls from the older adults Sound Life cohort (Ages 55-65, n = 31) were selected by a randomized sampling approach to univariately control for Sex, Race, Ethnicity, Age, and CMV status with the VRd cohort (n=17). The Wilcoxon Rank Sum test was used to test for continuous covariates and the Fisher’s Exact Test for categorical covariates. The randomized selection was repeated 7000 times to minimize confounding between covariates and cohorts, and the final cohort was selected as the set of controls that were most similar to the NDMM subjects. One additional subject was selected at random from the **younger adults** Sound Life cohort (Ages 25-35) to match the youngest age in the NDMM cohort, for a total of 32 healthy subjects.

#### Commercial source (BMMC Samples)

Frozen BMMCs from healthy patients were purchased from BIOIVT or StemCell technologies. Donors were purchased from the currently available catalog and no donors were collected prospectively for this study. All donors were prescreened for HIV-1 and 2, hepatitis B and hepatitis C as is standard. MNCs were then isolated according to standard isolation protocols; in brief; BMAs from self-reported healthy patients were collected into Na Hep or ACD A tubes and MNCs were isolated by layering over a density gradient, then MNCs were collected, washed and cryopreserved. All cells were stored in the gaseous phase of liquid nitrogen prior to purchase. Cells were purchased cryopreserved in CryoStor CS10, shipped on dry ice and stored in the gaseous phase of liquid nitrogen until use.

To obtain healthy BMIF, a fresh BMA from a healthy donor was purchased from Discovery Life Sciences (formerly AllCells) and shipped overnight at ambient room temperature. The BMA was diluted 1:1 in PBS + 2% FBS and processed over a density gradient to isolate BMMCs. The supernatant was collected, pooled and then centrifuged at 480xg for five minutes at room temperature. BMIF samples were aliquoted into 50 µL aliquots and stored at -80°C. Prior to Olink processing, samples were thawed and refrozen to match the freeze-thaw cycle of the on-study samples.

### Cell thaw

All cells were thawed per the protocol described in^79^. Briefly frozen mononuclear cells were removed from liquid nitrogen and immediately thawed in a 37°C water bath. Cells were diluted dropwise into 40 mL AIM V media (Thermo Fisher Scientific) pre-warmed to 37°C. Cells were pelleted at 400 x g for 5-10 min at 4-10°C, resuspended in 5 mL cold AIM V media, and counted using a Cellometer Spectrum or Cellaca. After adding 30 mL cold AIM V media, cells were re-pelleted and resuspended to the appropriate concentrations for scRNAseq or flow cytometry.

### 10x Genomics scRNA-seq v3 chemistry (PBMC samples)

Single-cell sequencing was performed on thawed PBMC samples using 10x Genomics Chromium 3′ v3.1 chemistry as previously described^23,79^. First, PBMCs were stained for oligonucleotide-tagged antibodies (HTO) and overloaded on Chip G wells (10x Genomics, 20000177). cDNA amplification was performed with HTO additive primer spike-in and then HTO and GEX products were separated using SPRI-Select (Beckman Coulter, B23319) bead-based cleanup before carrying forward into separate library indexing reactions. Libraries were sequenced using a NovaSeq X 25B 300 cycle flow cell or NovaSeq S4 200 cycle flow cell, depending on total read requirements at Northwest Genomic Center at the University of Washington (https://nwgc.gs.washington.edu/). Downstream sequencing data were computationally resolved and quality-checked using in-house pipelines.

### 10x CITE-seq (BMMC samples)

Up to one million cells from live BMMC samples were blocked with Human TruStain FcX Blocking Buffer (BioLegend, PN-422302) following manufacturer’s recommended usage. Cells were then stained and incubated for thirty minutes with oligo-tagged antibodies (HTOs) and a custom pool of DNA-barcoded antibodies (ADTs) (BioLegend, various PN). ADT-stained cells were washed to remove unbound antibodies and pooled for FACS sorting.

To remove dead cells and debris, cells were resuspended in 300 µL DPBS, Fixable Viability Stain 510 (BD) added and incubated for 30 min on ice protected from light. 10 mL sort staining buffer (AIM V medium plus 25 mM HEPES) was added and tubes centrifuged at 400g for 10 mins (4°C). Cells were adjusted to 10 million cells/mL in a sort staining buffer and passed through a 35 µm filter. Cells were sorted on a BD FACSAria Fusion cell sorter. A standard viable cell gating scheme was employed; FSC-A vs. SSC-A (to exclude sub-cellular debris), two FSC-A doublet exclusion gates (FSC-A vs. FSC-W followed by FSC-H), followed by a dead cell exclusion gate (BV510 LIVE/DEAD negative). An aliquot of each post-sort population was used to collect 10,000 events to assess post-sort purity.

After dead cell depletion via FACS sorting, cells were partitioned and barcoded using the Chromium Controller (10x Genomics) and the Chromium Next GEM Single Cell 3′ GEM, Library & Gel Bead Kit v3.1 (10x Genomics, PN-1000121). Up to 24 10x wells were loaded with total cells loaded per well ranging from 26,000 to 32,000 cells. At cDNA amplification, HTO-cDNA additive primer and ADT-cDNA additive primer were spiked into the cDNA amplification master mix and cDNA was amplified for 11 cycles following 10x Genomics cycling guidelines. Following cDNA amplification, HTO/ADT and cDNA products were separated using a 0.6X SPRI-Select (Beckman Coulter, PN B23319) bead-based cleanup as described in^80^, before carrying forward into separate library indexing reactions^79^. HTO and ADT libraries were amplified an additional 10 or 12 cycles with custom single-end index primers. Gene expression libraries were prepared in accordance with Chromium Next GEM Single Cell 3ʹ Reagent Kits v3.1 (CG000204, Rev D). Libraries were sequenced on a NovaSeq S4 200 cycle flow cell or NovaSeq X 25B 100 cycle flow cell at Northwest Genomic Center at the University of Washington (https://nwgc.gs.washington.edu/) following the manufacturer’s recommended sequencing guidelines (Read 1: 28cycles, Read 2: 91 cycles, index 1:8 cycles) targeting a minimum of 30,000 reads per cell for gene expression, 2,500 reads per cell for HTO, and 5,000 reads per cell for ADT libraries. Samples were then computationally resolved and quality-checked using in-house pipelines.

### Influenza-specific IgG serology assays

The Meso Scale DIscovery (MSD) Influenza Prototype 7-plex Serology Assay was performed as described in ^23^. Measures for IgG antibodies in human plasma specific for Influenza vaccine hemagglutinin antigens including A/Brisbane, A/Hong Kong, A/Michigan, A/Victoria, B/Colorado, B/Phuket and B/Washington. Samples were quantified in AU/mL and referenced against specific HA standard controls.

### SARS-CoV-2 IgG serology

The MSD 96-well V-Plex SARS-CoV-2 Panel 39 (IgG) Kit was run as per manufacturer’s instructions. All samples were diluted 1:100 in the kit provided diluent. Measures for antibodies in human plasma specific for SARS-CoV-2 N, SARS-CoV-2 Spike, SARS-CoV-2 Spike (B.1.617.2), SARS-CoV-2 Spike (BA.2.86), SARS-CoV-2 Spike (BA.5), SARS-CoV-2 Spike (JN.1 (v2)), SARS-CoV-2 Spike (KP.2), SARS-CoV-2 Spike (KP.3), SARS-CoV-2 Spike (LB.1), SARS-CoV-2 Spike (XBB.1.5 (v2)). Samples were quantified in AU/mL and referenced against specific HA standard controls.

### Olink Explore

The Olink Explore 1536 platform (Olink, Thermo Fisher Scientific) was used to measure the relative expression of 1,472 protein analytes in plasma samples (50 µL), and the Olink Explore 3072 platform was used to measure relative expression of 2,943 protein analytes in bone-marrow interstitial fluid (BMIF) samples (50 µL) at diagnosis and longitudinally across therapy. Olink data was normalized as per manufacturer’s instructions and as described in ^23^. Final normalized relative protein quantities were reported as log2-normalized protein expression (NPX) values.

### HCMV Serology

Serology testing for Human Cytomegalovirus (HCMV) was performed by the University of Washington’s Clinical Virology Laboratory in the Department of Laboratory Medicine (https://depts.washington.edu/uwviro/) as described in^23^. Patients were reported as CMV positive or negative assessed from CMV Ab Screen Index values.

### RISS stage and therapeutic responses

Patient response assessments were performed by the treating clinician at the time points in which the patient has a bone marrow sample collected (Pre-treatment, End Induction, Post-Transplant 90 days, 1-year Post-Transplant, and 2-years Post-Transplant). A combination of the IWMG response criteria ^2,9^ and data from pathology reports (% of abnormal plasma cells and cells via flow), imaging reports (e.g. PET CT), and disease marker labs (e.g. SPEP with IFE and UPEP) is used to assess individual patient response to therapy.

### Complete Blood Count (CBC), Serum Chemistry, Protein, Inflammatory, and Lipid Assessments

Complete blood count (CBC) and serum chemistry panels were collected as part of routine clinical evaluation. CBC parameters included both absolute and relative counts of major leukocyte subsets and erythroid indices: basophils, eosinophils, lymphocytes, monocytes, neutrophils, and immature granulocytes (reported as counts and percentages), as well as hematocrit, hemoglobin, red blood cell count (RBC), white blood cell count (WBC), platelet count, mean corpuscular volume (MCV), mean corpuscular hemoglobin (MCH), mean corpuscular hemoglobin concentration (MCHC), and red cell distribution width (RDW and RDW-CV).

Serum chemistry analyses included markers of liver function (ALT, AST, alkaline phosphatase, total bilirubin, albumin, and total protein), renal function (creatinine, blood urea nitrogen [BUN]), electrolyte balance (sodium, potassium, chloride, CO₂, anion gap, calcium, magnesium, phosphate), glucose metabolism (glucose), and systemic inflammation or cell turnover (lactate dehydrogenase [LDH]). All laboratory testing was performed using standard clinical analyzers in a CLIA-certified laboratory.

To complement cellular and molecular analyses, additional serum-based biomarkers were measured. Immunoglobulin profiling included quantification of IgA, IgG, and IgM, as well as serum immunofixation to detect monoclonal protein species. Free light chain (FLC) analysis included kappa and lambda FLC concentrations and the kappa/lambda ratio. Additional monoclonal protein characterization included identification and quantification of monoclonal spikes.

Inflammatory markers included erythrocyte sedimentation rate (ESR) and high-sensitivity C-reactive protein (hsCRP), serving as systemic indicators of acute and chronic inflammation. Lipid panels included high-density lipoprotein (HDL), low-density lipoprotein (LDL), total cholesterol, non-HDL cholesterol, triglycerides, and the cholesterol:HDL ratio, assessed as part of cardiometabolic risk profiling.

Serum protein electrophoresis (SPE) measures included total protein, albumin, and globulin fractions (alpha-1, alpha-2, beta, and gamma). Beta-2 microglobulin (β2M) was also measured as a marker of tumor burden and renal function.

### Clinical Cytogenetics

Cytogenetic data were derived from fluorescence in situ hybridization (FISH) or karyotype analyses performed on bone marrow aspirates at diagnosis or follow-up. Assessed abnormalities included common myeloma-associated chromosomal changes: deletion of 17p (cyto.17p_loss), gain of 1q (cyto.1q_gain), and deletion of 1p (cyto.1p_loss). Structural rearrangements involving immunoglobulin heavy chain (IgH) loci were also recorded, including translocations t(11;14), t(14;16), t(14;20), and t(4;14). A composite high-risk cytogenetic designation (cyto.high_risk) was assigned based on the presence of one or more high-risk lesions, following IMWG or institutional criteria. Karyotype (cyto.karyotype) derived from clinical FISH, is described as abnormal or normal in a subset of patients.

### HLA, KIR, and FcGR Genotyping

Coded PBMC samples were submitted for high resolution genotyping of human leukocyte antigen (HLA), Killer cell immunoglobulin-like Receptor (KIR), and Fc gamma receptor alleles by <u>Scisco</u> <u>Genetics</u>, Inc., Seattle, WA using short-read next generation sequencing assays. The ScisGo-HLA-v6 technology was utilized to identify up to 11 different HLA-class I and -class II loci based on manufacturer’s instructions. The ScisGo-KIR-v3 system was run per the manufacturer’s instructions as previously described ^81^.

### Flow Cytometry Data Analysis

Flow cytometry staining was performed as described in ^26^. PBMCs were thawed, split into four equal aliquots and stained with one of four panels, PS1, PT1, PB1 or PM1. BMMCs were thawed, split into two equal aliquots and stained with one of two panels; BMS1 or BMM1. Complete panel details including antibody, clone, vendor and titre were listed in the reagents table shown. All flow cytometry data was analyzed using CellEngine. PBMCs were defined as described in ^26^ with the exception of Tfh cells in the PT1 panel and Plasma cells in the PB1 panels. Tfh cells were defined as CD185⁺, CD279(PD-1)⁺ within the non Treg population (CD4⁺, CD25-) and plasma cells were defined as CD19⁺, CD20-, CD27⁺, CD319⁺ due to the CD38 degradation seen in patients in the DVRd cohort.

BMMC core immune cell populations, T, B, NK and myeloid subsets, were defined as similarly as possible to the PBMCs so that most major cell types were captured in both cell types. Additionally HSCs were defined as CD34⁺, lin-, CD38-, CD45RA-.

Representative flow plots shown were generated in Cell Engine by concatenating all cells from each timepoint together and then downsampling to show the same representative cell numbers.

Gated cell type frequencies exported from CellEngine for each panel were combined into one dataset prior to processing. Cell type nomenclature was standardized across panels by consolidating variant labels (e.g., "B cells", "B Cells", "Total B Cells") into uniform identifiers. Samples were filtered to include only Multiple Myeloma-relevant timepoints: Healthy controls, Pre-Treatment (PreTx), Post Induction 2-Cycles (PI2C), End Induction (EI), Post Transplant 60 Days (ASCT60d), Post Transplant 90 Days (ASCT90d), Post Transplant 1 year (ASCT1y), and Post Transplant 2 years (ASCT2y). For subjects missing the 60-day post-transplant timepoint but with available 90-day data, the 90-day measurement was substituted as a proxy for the 60-day timepoint.

To derive estimated absolute cell counts from flow cytometry data, we normalized each cell population using the patient’s clinical Absolute Lymphocyte Count (ALC). For each sample, total lymphocyte counts (T cells + B cells + NK cells) were first calculated from the flow cytometry data. The estimated absolute count for each cell type was then computed as:

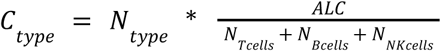

Where *C_type_* represents the estimated absolute count for a given cell type, *N_*type*_* is the event count for that cell type from flow cytometry, *ALC* is the clinical absolute lymphocyte count (cells/μL), and the denominator represents the total lymphocyte count from flow cytometry.

Following ALC normalization, total B, T, and NK cell populations were removed from subsequent analyses to focus on immune cell subsets.

For samples with duplicate measurements at the same subject-visit-celltype combination, count values were averaged to generate a single representative measurement per group.

Cell type frequencies were calculated separately for each sample by dividing individual cell type counts by the sum of all measured cell types within that sample, ensuring frequencies summed to 1.0 per sample. Both raw frequencies (based on event counts) and ALC-normalized frequencies (based on estimated absolute counts) were computed. To account for the compositional nature of the data, we applied a centered log-ratio (CLR) transformation. For each sample, the CLR value for cell type **i** was calculated as:

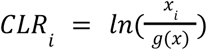

Where x_i_ is the frequency of cell type i and g(x) is the geometric mean of all cell type frequencies in that sample. Each flow cytometry panel (PS1, PT1, PB1, PM1) was processed independently through the ALC normalization, duplicate handling, frequency calculation, and CLR transformation pipeline to preserve panel-specific cell populations. The survey panel (PS1) was processed identically with one modification: B cells in this panel were labeled as "Partial B Cells" as this measurement excludes plasma cells. Progenitor cells (circulating CD34⁺) were measured exclusively in the survey panel and were subsequently merged with the processed data from the other panels to generate a unified dataset containing all cell types.

### Bone Marrow scRNAseq QC and Cell Labelling

Bone marrow mononuclear cell (BMMC) samples were processed using Scanpy. Doublets were identified using Scrublet with default parameters. Cell type annotations were transferred using CellTypist from three reference datasets: (1) healthy PBMC reference ^23^, (2) bone marrow dataset ^82^, and (3) internal 5’ Flex kit bone marrow dataset available at https://doi.org/10.57785/va48-4c35. All cells were normalized using counts per million and log1p-transformed prior to annotation.

### Isolation and processing of malignant plasma cells

To preserve patient-specific tumor heterogeneity, malignant plasma cells were isolated separately from non-malignant immune cells. Plasma cells were identified using expression of canonical markers (*CD38*, *TNFRSF17*, *SDC1*, *PRDM1*, *JCHAIN*) and immunoglobulin genes (*IGHG1-4*, *IGHA1-2*, *IGHD*, *IGHM*, *IGHE*, *IGLC1/2/3/6/7*, *IGKC*) across unsupervised Leiden clusters. A second iteration of clustering and marker-based filtering removed residual doublets and mislabeled cells. For downstream analysis of plasma cells, mitochondrial (MT-), ribosomal (RPS, RPL), hemoglobin (HB), and immunoglobulin genes were masked to minimize technical artifacts. Plasma cell barcodes were excluded from subsequent non-malignant cell processing.

### Quality control and processing of non-malignant bone marrow cells

The following steps were performed for all healthy bone marrow samples and MM samples excluding malignant cells. Quality control metrics were calculated including mitochondrial, ribosomal, and hemoglobin gene content. Outlier cells were identified using median absolute deviation (MAD) thresholds: 7 MADs for total counts, genes detected, and top 20 gene expression; >6% for mitochondrial content; and >0.35 for doublet score. Following quality filtering, cells were normalized to 10,000 counts per cell and log1p-transformed. Highly variable genes were identified (min_mean=0.0125, max_mean=3, min_disp=0.25), followed by scaling (max_value=10) and PCA. Batch correction was performed using Harmony when indicated. Neighborhood graphs were constructed using 50 neighbors and 20 principal components, followed by TSNE dimensionality reduction (n_pcs=20), UMAP dimensionality reduction (min_dist=0.45, random_state=0, n_components=2) and Leiden clustering (resolution=2.0). This function is available for reuse at (https://github.com/aifimmunology/2025-mm-project/blob/development/rna-base-functions/process_ <u>scrna_data.py</u>)

### BMMC Cell Type Annotation

Major lineages were defined using canonical markers: T cells (*CD3D*, *CD3E*, *CD2*, *TRAC*, *IL7R*, *CD4*, *CD8A*); NK cells (*NKG7*, *GNLY*, *KLRD1*, *KLRF1*, *FCGR3A*, *PRF1*); B cells (*CD19*, *CD79A*, *MS4A1*, *CD22*); monocytes (*CD14*, *LYZ*, *S100A8*, *S100A9*); dendritic cells (*CST3*, *CLEC9A*, *CD1C*, *FCER1A*, *LILRA4*); mesenchymal stromal cells (*LEPR*, *ENG*, *THY1*, *CXCL12*); and progenitors (*CD34*, *KIT*, *MKI67*). Fine-grained cell type assignments were manually refined based on marker gene expression patterns within these major lineages. Residual doublets and outliers identified during manual curation were removed. This rigorous annotation process yielded a hierarchical classification system with 64 unique cell types at the finest resolution (Level 3), which were grouped into 39 cell types at intermediate granularity (Level 2) and 7 cell types at low granularity (Level 1). The final dataset has 1171965 BMMC cells.

### Peripheral Blood scRNA-seq QC and Cell Labelling

Quality control metrics were calculated for mitochondrial gene content and genes detected per cell. Cells were filtered using the following thresholds: <10% mitochondrial content, 200-5000 genes detected, and predicted doublet status removed. A secondary round of doublet detection was performed on the filtered dataset, and additional doublets were removed to ensure high-quality singlets. Cell type annotations were transferred from the Allen Institute for Immunology healthy PBMC reference atlas ^23^ using CellTypist. All cells were normalized to counts per million and log1p-transformed prior to annotation as recommended by CellTypist. Transferred labels included fine-grained annotations at three hierarchical levels (L1, L2, L3). Annotations were manually reviewed based on canonical marker expression, and ambiguous or low-confidence labels were removed. Intermediate cell types that fail automated label transfer, like intermediate monocytes were defined manually based on marker expression. Cells annotated as doublets or low-quality cells during manual curation were excluded from downstream analyses. The healthy PBMC reference remains as defined in ^23^. The final dataset has 5413474 PBMC cells.

### Single-cell Data Pseudobulking

We aggregated single-cell data by summing counts within each sample, reducing noise and dropouts in a process called pseudobulking according to the formula:

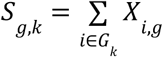

where *S*_*g*,*k*_ = summed count of gene *g* in group *k*,

*X*_*i*,*g*_ = raw count of gene *g* in cell *i*, and

*G*_*k*_ = set of all cells belonging to group *k*.

Cell types were filtered to a minimum of 10 cells per cell type, and samples were filtered for a minimum of 1000 cells per sample.

### Pathway Enrichment

Genes were ranked according to their significance (-log10(p-value)) and direction from DESeq2, and analysed using the FGSEA R package (cite). Databases included were MSigDB_Hallmark_2020, Reactome_Pathways_2024, ENCODE_and_ChEA_Consensus_TFs_from_ChIP-X, KEGG_2021_Human. Significant pathways with an adjusted p-value <= 0.05 were visualized.

### CNV Analysis

From the scRNA-Seq analysis, data from labeled BMMC malignant and non-malignant plasma cells, along with publicly available data from healthy plasma cells was selected. The transcripts were normalized, followed by scaling, variable feature selection, PCA, and UMAP embedding as described in Scanpy (cite). A healthy bone marrow external atlas dataset from researchers at the University of Pennsylvania was subsetted down to plasma cells (n=16K) (http://cell.com/cell/fulltext/S0092-8674(24)00408-2) and merged with our NDMM plasma cells and harmonized on data source. Chromosome locations for each gene were added to run InferCNV. InferCNV ran with reference keys Healthy vs NDMM Plasma and reference group being the healthy external atlas dataset (window size = 200). The CNV object was processed with PCA, UMAP embedding, leiden clustering and nearest neighbor clustering and CNV score was calculated using the inferCNV method^83^.

### Olink Pathway Analysis

Pathway enrichment analysis of DEPs across contrasts was conducted using the Over-Representation Analysis (ORA) method implemented in WebGestaltR. Databases included the Encode, KEGG, MsigDB, and Reactome pathway databases accessed through EnrichR. Proteins were clustered according to fold change across pairs of timepoints to identify clusters of similar longitudinal trajectories.

### Statistical analysis

#### Hypothesis testing for Gene Expression Data

Statistical analyses were performed in the R Statistical Software (v4.3.3; R Core Team 2024).

To identify differentially expressed genes (DEGs), genes had to be:

- expressed in >=10% of the cell types were included, and
- be expressed in a minimum of 5 samples per condition and cell type.

Differential genes analysis was conducted in pseudobulk using the DESeq2 package in R ^84^ as follows:

- NDMM Paired Analysis: For evaluating pseudobulk gene expression changes between two clinical visits, a paired analysis was done following DESeq2’s recommendations of regressing subject-specific effects using the following formula *GEX* ∼ *Subject* + *Visit*.
- NDMM Comparisons vs healthy cohorts: We tested pseudobulk expression differences against healthy matched controls adjusting for age, sex, and CMV status using the following formula: *GEX* ∼ *Cohort* + *Sex* + *Age* + *CMV*.

Benjamini-Hochberg p-value correction was applied to control the false discovery rate (FDR), with significant differences in composition defined by FDR < 0.05. Differential genes were defined by an adjusted p<= 0.05.

#### Hypothesis Testing for Compositional and Proteomic Data

Statistical differences between healthy and cancer patients (VRd/DVRd) for each timepoint were calculated using an unpaired Mann-Whitney U test. Statistical differences between timepoints within the same treatment group (Healthy, VRd, or DVRd) were calculated using a paired Wilcoxon Signed-Rank test. Benjamini-Hochberg p-value correction was applied to control the false discovery rate (FDR), with significant differences in composition defined by FDR < 0.05.

### B cell isotype classification

Normalized expression of nine IgH constant genes (*IGHM*, *IGHD*, *IGHG1-4*, *IGHA1-2*, and *IGHE*) were used to classify B cells by their isotype. If cells were positive for either *IGHM* or *IGHD*, or both *IGHM* and *IGHD*, they received isotype assignment of *IGHM*, *IGHD*, or *IGHMD*. Otherwise, the cell received the isotype of the IGH genes for which they are positive. In case a cell is positive for multiple IGH genes, they are assigned the isotype for the positive gene closest to the variable region in genomic DNA, which follows the order: *IGHG3*, *IGHG1*, *IGHA1*, *IGHG2*, *IGHG4*, *IGHE*, *IGHA2*. In cases of no expression of any IGH genes, the cell was classified as “undetermined”.

