## Supplemental Figures for "Myeloma and therapy reshape the bone marrow niche to durably constrain immune reconstitution and vaccine responsiveness"

### Supplemental figures and legends

Supplemental Figure 1 (Relevant to Figure 2)

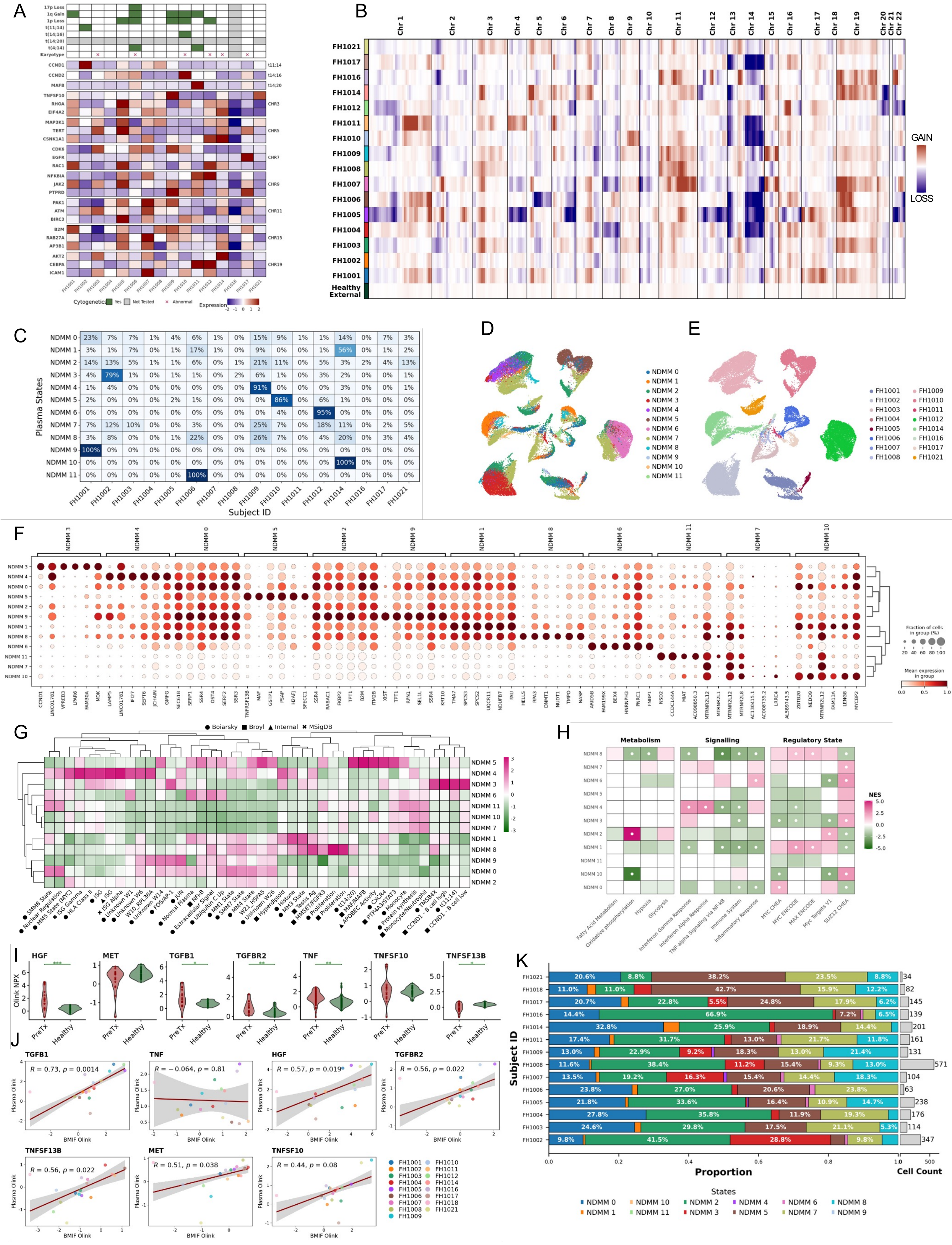

**Supplemental Figure 1 (related to Figure 2). Genetic context for NDMM tumor states using cytogenetics, expression, and inferred copy number.**

**(A)** Integrated cytogenetic abnormalities and tumor gene expression across the NDMM cohort, with cytogenetic calls indicated (detected, not tested, abnormal karyotype) and z-scored expression of representative genes grouped by translocation partner or chromosomal region. **(B)** Inferred copy number variation from residual expression intensities relative to a healthy baseline, with gains and losses indicated by the color scale. **(C)** Per-donor contribution of malignant plasma cells to each Leiden tumor state. **(D)** UMAP of NDMM plasma cells at PreTx colored by harmonized Leiden states (NDMM_0-NDMM_11). **(E)** UMAP of NDMM plasma cells colored by patient. **(F)** Marker-gene dot plot for Leiden tumor states, with gene groups labeled by the associated state. **(G)** Clustered heatmap of published MM gene signature scores across Leiden tumor states, with signatures ordered by hierarchical clustering. **(H)** FGSEA enrichment (NES) using DEGs from each Leiden state versus healthy donor plasma cells across selected pathways. **(I)** Differential abundance of selected tumor-associated plasma proteins at PreTx versus healthy donors, with significance indicated as shown (AdjP < 0.05, AdjP < 0.01, AdjP < 0.001). **(J)** Spearman correlations of tumor-associated proteins between matched BMIF and plasma samples at diagnosis (NPX). **(K)** Per-donor plasma cell state composition after end of induction, with total tumor cell counts per donor. Abbreviations: newly diagnosed multiple myeloma (NDMM), uniform manifold approximation and projection (UMAP), gene set enrichment analysis (FGSEA), normalized enrichment score (NES), differential expressed genes (DEGs), copy number variation (CNV), bone marrow interstitial fluid (BMIF), normalized protein expression (NPX), pre-treatment (PreTx), end of induction (EI).

Supplemental Figure 2 (Relevant to Figure 3)

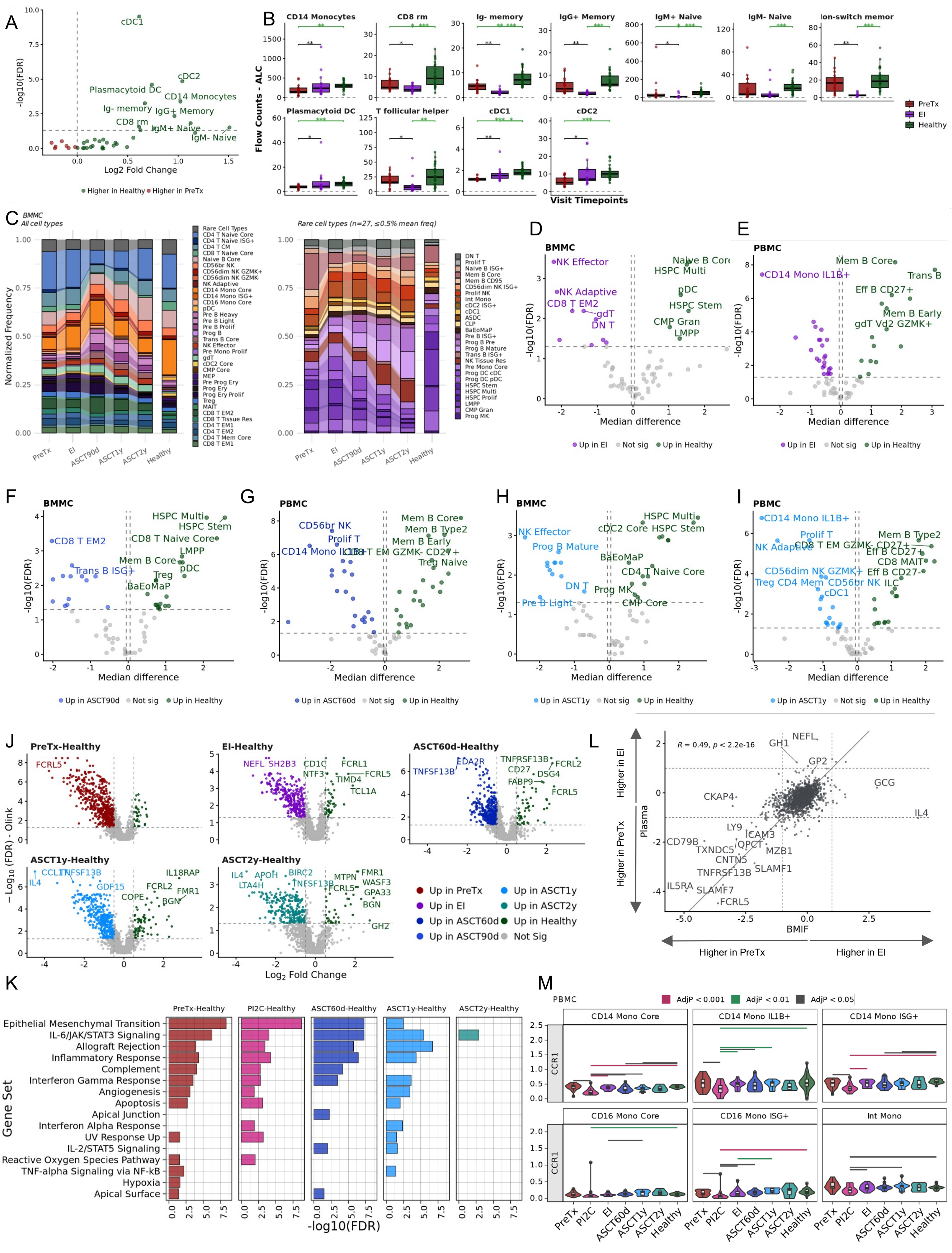

**Supplemental Figure 2 (related to Figure 3). Peripheral and marrow immune composition with matched plasma and BMIF proteomic shifts across therapy.**

**(A)** Volcano plot of flow cytometry PBMC composition at PreTx versus healthy donors, with direction of change indicated by the color key. **(B)** Flow cytometry PBMC subset absolute counts in PreTx NDMM (VrD subjects) versus healthy donors (box and whisker plots), with significance annotated (AdjP * < 0.05, AdjP ** < 0.01, AdjP *** < 0.001). **(C)** scRNA-seq BMMC compositional alluvial plot by normalized frequency across timepoints with rare populations collapsed in the main view and shown in a rare-only view, with healthy donors shown for reference. **(D-I)** Volcano plots of scRNA-seq compositional differences versus healthy donors at each timepoint, summarized as median CLR difference (x-axis) and FDR (y-axis), shown for BMMCs (**D, F, H**) and PBMCs (**E, G, I**). **(J)** Volcano plots of plasma protein differential abundance at each NDMM timepoint versus healthy donors (log2FC on x-axis). **(K)** MSigDB pathway enrichment from differentially abundant proteins at each NDMM timepoint versus healthy donors. **(L)** Spearman correlation of protein log2FC between BMIF and plasma for the PreTx versus EI contrast. **(M)** Pseudobulk CCR1 expression in PBMC monocyte populations, with significance encoded by the bracket color key (maroon, AdjP< 0.001; seagreen, AdjP < 0.01; gray, AdjP < 0.05. Abbreviations: pre-treatment (PreTx), end of induction (EI), peripheral blood mononuclear cells (PBMC), bone marrow mononuclear cells (BMMC), bone marrow interstitial fluid (BMIF), centered log-ratio (CLR), false discovery rate (FDR), differentially expressed proteins (DEPs), Molecular Signatures Database (MSigDB), log2 fold change (log2FC).

Supplemental Figure 3 (Relevant to Figure 3)

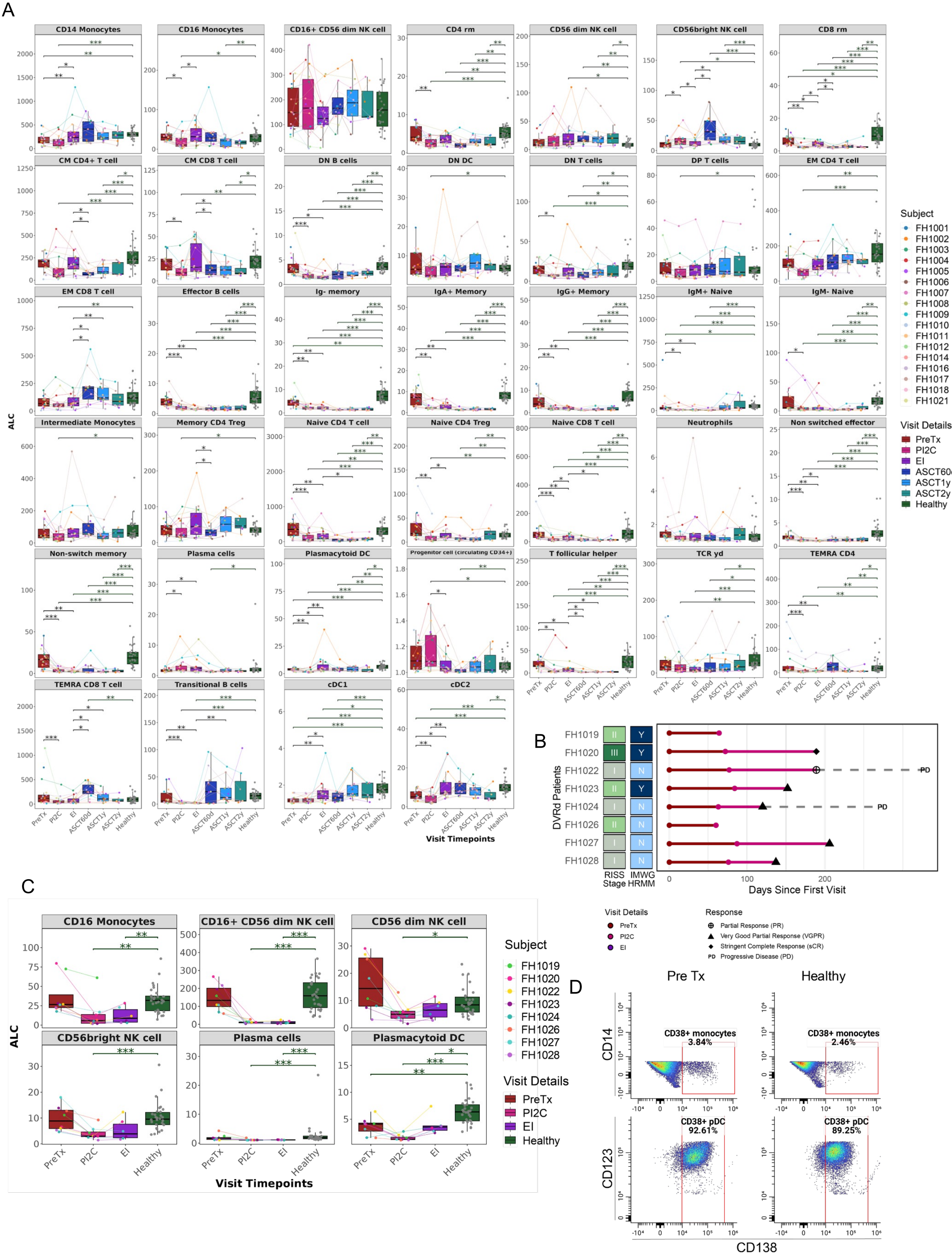

**Supplemental Figure 3 (related to Figure 3). Flow cytometry validation of PBMC immune composition changes across induction cohorts.**

**(A)** Flow cytometry PBMC subset frequencies across visits in the VRd cohort, with significance bars indicating comparisons versus healthy (black), within induction timepoints (red), and post-induction comparisons (purple), with p-value thresholds as shown (AdjP * < 0.05, AdjP ** < 0.01, AdjP *** < 0.001). **(B)** Sample manifest and clinician-recorded responses for NDMM participants receiving DVRd (n = 8). **(C)** Flow cytometry PBMC subset frequencies across visits in the DVRd cohort with the same comparison scheme and significance thresholds as in **(A)**. **(D)** Concatenated flow cytometry showing CD38 expression in monocyte and dendritic cell populations across VRd timepoints, shown in VRd (rather than DVRd) to avoid daratumumab-associated depletion and masking of CD38+ populations that can confound apparent composition. Abbreviations: newly diagnosed multiple myeloma (NDMM), peripheral blood mononuclear cells (PBMC), bortezomib/lenalidomide/dexamethasone (VRd), daratumumab plus VRd (DVRd), dendritic cell (DC).

Supplemental Figure 4 (Relevant to Figure 4)

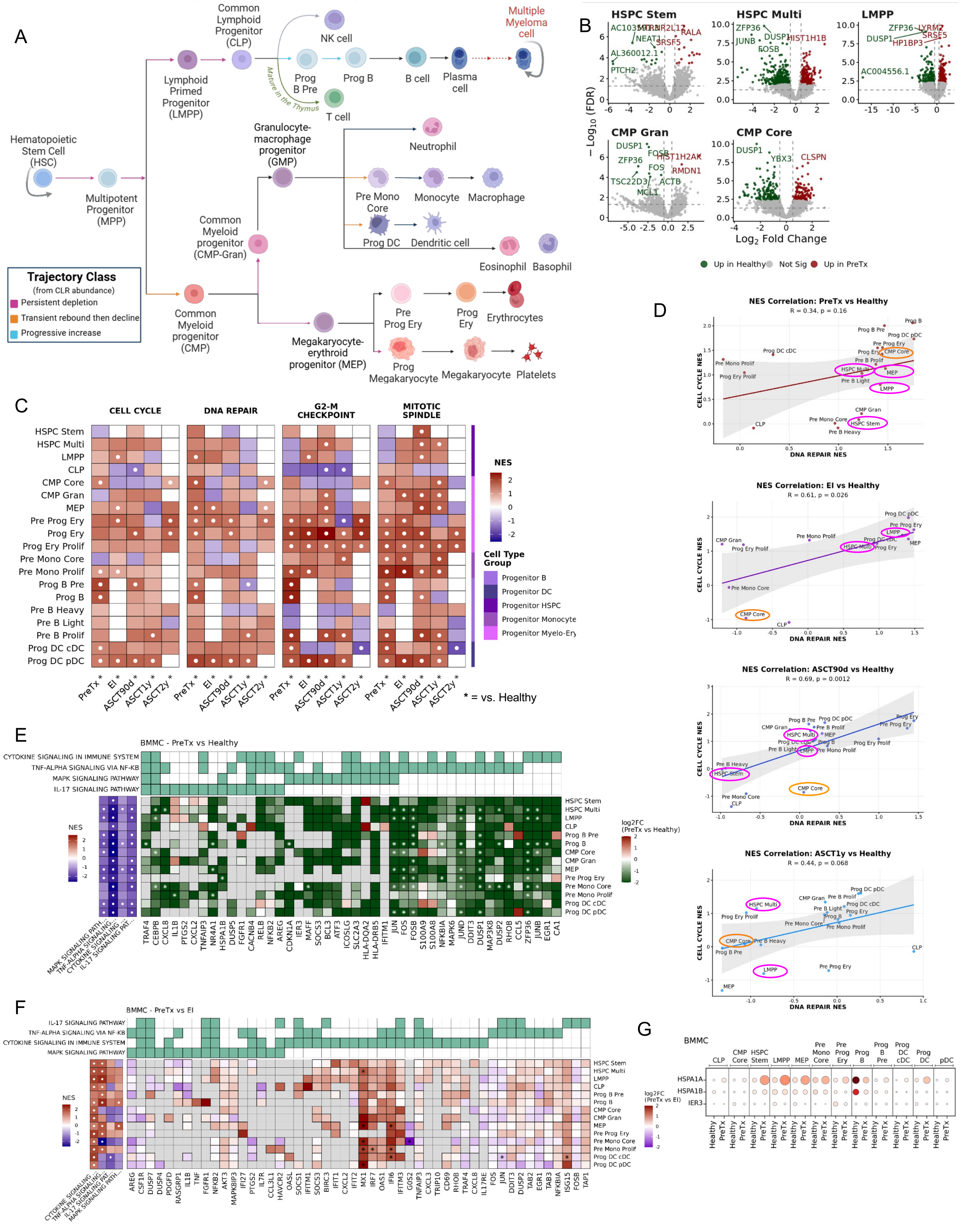

**Supplemental Figure 4 (related to Figure 4). Hematopoietic progenitor trajectory remodeling and pathways across treatment timepoints.**

**(A)** Schematic of hematopoietic differentiation with progenitor and immune lineages arranged along canonical trajectories, with colored arrows summarizing subset dynamics across therapy based on CLR abundance versus healthy (persistent depletion, transient rebound then decline, progressive increase, as labeled). **(B)** Volcano plot of differential expression in progenitor populations comparing NDMM PreTx versus healthy donors (log2FC on x-axis, FDR on y-axis), with direction and significance indicated by the color key. **(C)** Pathway enrichment (NES) across progenitor cell types at NDMM timepoints relative to healthy controls. **(D)** Spearman correlation of DNA repair and cell cycle NES across progenitor cell types at successive timepoints relative to healthy, with trend-lines and 95% confidence intervals overlaid. Ovals represent trajectory class from (A). **(E-F)** Pathway-level heatmaps and leading-edge gene heatmaps for the indicated contrasts (**E**), PreTx vs. Healthy and (**F**), PreTx vs EI, including pathway membership annotations for leading-edge genes. **(G)** Dot plot of heat shock and stress gene expression in PreTx and healthy progenitor populations. For FGSEA, white dots denote AdjP<0.05. Abbreviations: pre-treatment (PreTx), end of induction (EI), centered log-ratio (CLR), false discovery rate (FDR), normalized enrichment score (NES), log2 fold change (log2FC).

Supplemental Figure 5 (Relevant to Figure 5)

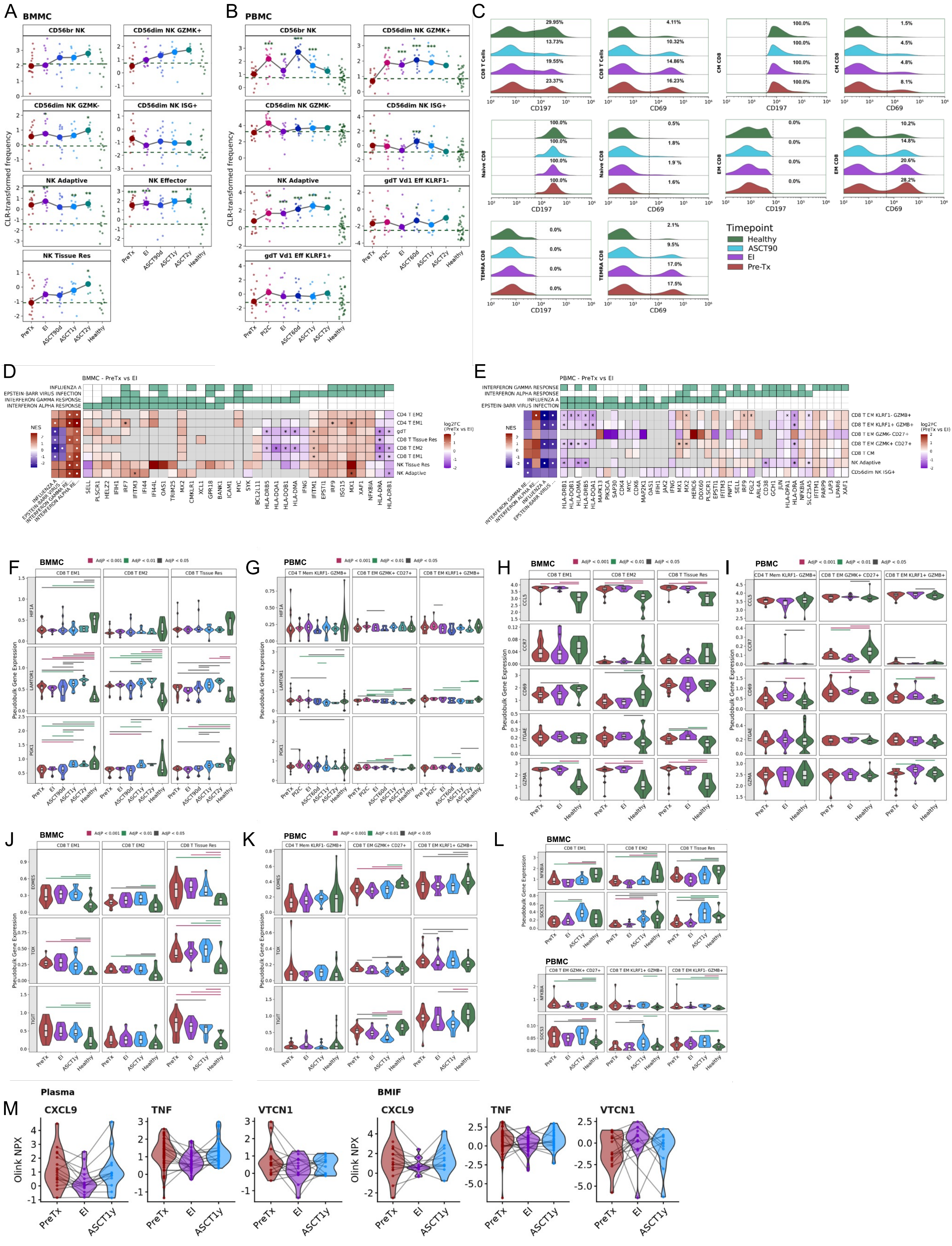

**Supplemental Figure 5 (related to Figure 5). Supporting analyses for cytotoxic lymphocyte dynamics including NK kinetics and immune pathway leading-edge structure.**

**(A-B)** CLR-transformed frequencies of NK cell subsets across timepoints relative to healthy controls in BMMCs **(A)** and PBMCs **(B)**, with significance annotated (AdjP * < 0.05, AdjP ** < 0.01, AdjP *** < 0.001). **(C)** Concatenated flow cytometry histograms of CCR7 (CD197) and CD69 expression in BMMC immune subsets across timepoints (color key as shown). **(D-E)** Heatmaps of pathway-level changes and leading-edge genes for the PreTx versus EI contrast in BMMCs **(D)** and PBMCs **(E)**, with pathway membership annotations. For FGSEA, white and black dots denote AdjP<0.05. **(F-L)** Pseudobulk expression across cytotoxic T cell subsets in BMMCs (**F, H, J, L**) and PBMCs (**G, I, K, L**), with significance encoded by the bracket color key (maroon, AdjP < 0.001; gray, AdjP * < 0.05; seagreen, AdjP * < 0.01). **(M)** BMIF (right) and plasma (left) proteomics across NDMM treatment timepoints. Abbreviations: natural killer (NK), bone marrow mononuclear cells (BMMC), bone marrow interstitial fluid (BMIF), peripheral blood mononuclear cells (PBMC), pre-treatment (PreTx), end of induction (EI), centered log-ratio (CLR).

Supplemental Figure 6 (Relevant to Figure 5)

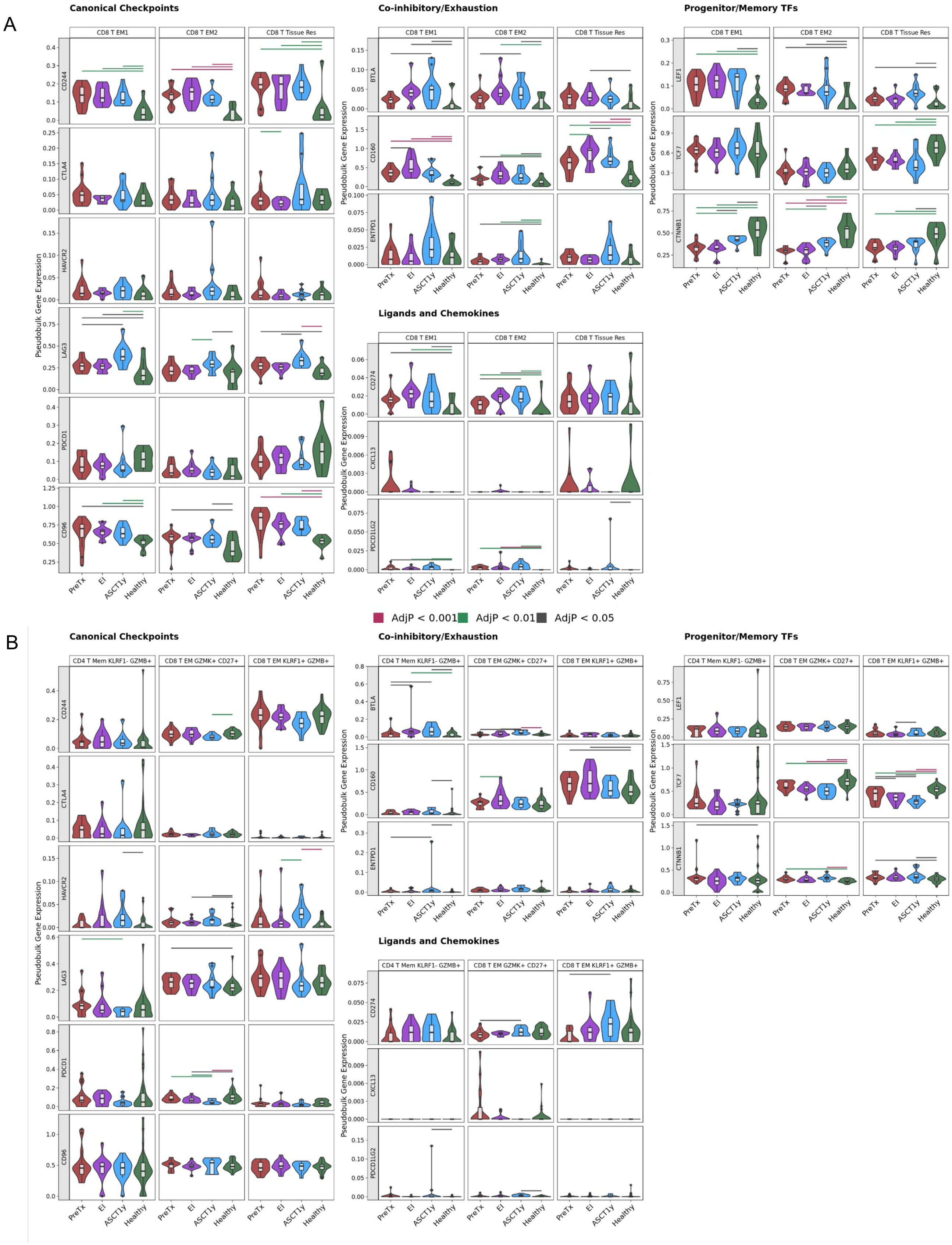

**Supplemental Figure 6 (related to Figure 5). Expanded pseudobulk marker panels for cytotoxic T cell regulatory gene sets in marrow and blood.**

**(A)** Pseudobulk expression panels across BMMC T cell subsets, grouped by the gene set category indicated above each panel, with significance encoded by bracket color (maroon, AdjP < 0.001; gray, AdjP < 0.05; seagreen, AdjP < 0.01). **(B)** Parallel pseudobulk expression panels across PBMC T cell subsets with the same gene set groupings and significance encoding. Abbreviations: bone marrow mononuclear cells (BMMC), peripheral blood mononuclear cells (PBMC).

Supplemental Figure 7 (Relevant to Figure 6 and Figure 7)

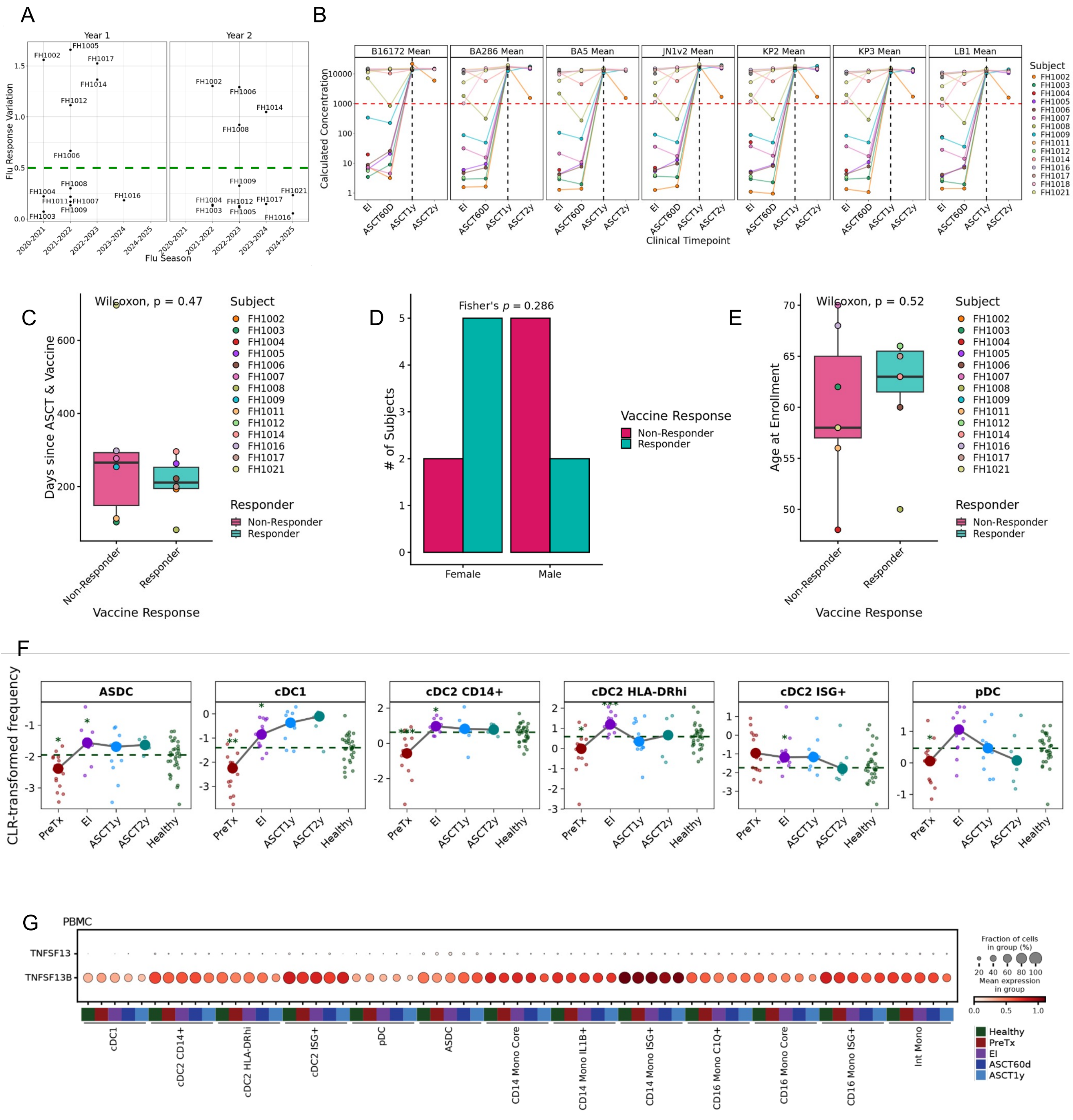

**Supplemental Figure 7 (related to Figures 6 and 7). Peripheral immune recovery and vaccine response analysis across DC and B cell compartments.**

**(A)** Flu response by season, with responder threshold indicated (green line). **(B)** Longitudinal SARS-CoV-2 vaccine responses across NDMM treatment timepoints (strains as labeled). **(C)** Vaccine response stratified by time from ASCT to vaccination, grouped by responder status. **(D)** Sex distribution across responder groups. **(E)** Age at enrollment stratified by responder group. **(C-E)** Nominal p-values are shown to test differences in flu response time to vaccine, sex, and age, respectively. **(F)** Longitudinal dynamics of selected progenitor populations across treatment timepoints relative to healthy controls. Significance annotated (AdjP * < 0.05, AdjP ** < 0.01, AdjP *** < 0.001). **(G)** Dot plots across myeloid subsets and timepoints with comparisons versus healthy controls. Abbreviations: autologous stem cell transplant (ASCT), newly diagnosed multiple myeloma (NDMM), dendritic cell (DC).
